# Greater leaf photosynthesis in the field by increasing mesophyll conductance via modified cell wall porosity and thickness in tobacco

**DOI:** 10.1101/2024.02.13.580201

**Authors:** Coralie E. Salesse-Smith, Edward B. Lochocki, Lynn Doran, Benjamin E. Haas, Samantha S. Stutz, Stephen P. Long

**Affiliations:** Carl R. Woese Institute for Genomic Biology, University of Illinois at Urbana-Champaign, 1206 W. Gregory Dr., Urbana, IL 61801, USA; Departments of Plant Biology and of Crop Sciences, University of Illinois at Urbana-Champaign, 505 South Goodwin Avenue, Urbana, IL 61801, USA

**Keywords:** Mesophyll conductance, Cell wall thickness, CO_2_ assimilation, Carbon isotope discrimination, *AtCGR3*, Pectin methyltransferase, water use efficiency, photosynthetic efficiency, Cell wall conductance, Effective porosity

## Abstract

Mesophyll conductance (*g*_m_) describes the ease with which CO_2_ passes from the sub-stomatal cavities of the leaf to the primary carboxylase of photosynthesis, Rubisco. Increasing *g*_m_ has been suggested as a means to engineer increases in photosynthesis by increasing [CO_2_] at Rubisco, inhibiting oxygenation and accelerating carboxylation. Here tobacco was transgenically up-regulated with Arabidopsis Cotton Golgi-related 3 (*CGR3*), a gene controlling methylesterification of pectin, as a strategy to increase CO_2_ diffusion across the cell wall and thereby increase *g*_m_. Across three independent events in tobacco strongly expressing *AtCGR3,* mesophyll cell wall thickness was decreased by 7-13%, wall porosity increased by 75%, and *g*_m_ measured by carbon isotope discrimination increased by 28%. Importantly, field-grown plants showed an average 8% increase in leaf photosynthetic CO_2_ uptake. Upregulating *CGR3* provides a new strategy for increasing *g*_m_ in dicotyledonous crops, leading to higher CO_2_ assimilation and a potential means to sustainable crop yield improvement.

---

Photosynthesis, the process of converting light energy and atmospheric CO_2_ into organic compounds, is directly or indirectly the source of all food. Improving photosynthetic efficiency has become a major research objective in order to feed an increasing global population, and to supplement ongoing crop breeding efforts^1,2^. A critical need is to achieve increases without the use of more land or water, given pressures on supply and diminished soil moisture under climate change^3–5^. One strategy with the potential to help meet this challenge is to use genetic engineering to increase photosynthetic efficiency of C_3_ plants via increased mesophyll conductance^6–8^. However, in order to test this, there is a need to gain a better understanding of mesophyll conductance and how manipulating it may affect photosynthesis and water use efficiency.

Mesophyll conductance (*g*_m_) measures the ease with which CO_2_ from the sub-stomatal cavities may diffuse to the chloroplast stroma, where it is fixed by Rubisco. Increasing *g*_m_ can increase photosynthetic capacity of C_3_ plants, and potentially crop yields, by increasing the concentration of CO_2_ around Rubisco^7,9^. This would decrease photorespiratory losses and accelerate carboxylation, without any additional cost in transpiration^7,8^. A combination of factors are considered to affect *g*_m_. These include gas phase diffusion from the inside of the stomata to exposed mesophyll cell walls and then liquid phase diffusion through the cell wall, plasma membrane, cytosol, chloroplast envelope and chloroplast stroma^10–13^.

Mesophyll conductance is influenced by several leaf anatomical properties^10^. Among these are the chloroplast surface area exposed to intercellular airspaces (*S*_c_), the mesophyll surface area exposed to intercellular airspaces (*S*_m_), and their ratio (*S*_c_/*S*_m_), the latter of which has been shown to be positively correlated with *g*_m_^14^. Mesophyll cell wall thickness (*T*_cw_) and porosity, as well as the permeability of the plasma membrane and chloroplast envelope to CO_2_, are also considered important properties affecting *g*_m_^15,16^. Both aquaporins and plastid surface area have been suggested to affect *g*_m_^6,17^. However, manipulation studies have produced mixed results^18–21^. Several modelling studies have suggested that the cell wall is one of the most prominent constraints on *g*_m_^12,22,23^. Cell wall conductance to CO_2_ (*g*_cw_) depends on its thickness, the tortuosity of the path of CO_2_ through the pores of the cell wall (*τ*), and the number of those pores (porosity *p*)^10^. Previous studies, including one on natural variation with leaf age in tobacco leaves, have reported that 1/*g*_m_ has a strong positive correlation with cell wall thickness, inferring that decreasing cell wall thickness is a means to increase *g*_m_ ^24,25^.

Cell wall formation is a complex process involving many genes and their protein products, so there are many potential options for altering cell wall thickness. Previous studies in *A. thaliana* have shown that overexpression of Cotton golgi related 3 (*AtCGR3*) or a functionally redundant gene *AtCGR2* increased the fraction of intercellular airspaces (*f*_ias_) and plant growth^26,27^. CGR3 is a pectin methyltransferase that catalyzes the methylesterfication of pectin in the cell wall^26^. Essentially, CGR3 adds methyl groups to pectin, serving to increase the extensibility of the cell wall, while affecting porosity ^27,28^. Pectin is one of the three main components of dicot primary cell walls, along with cellulose and hemicellulose. Increasing the ratio of pectin to cellulose and hemicellulose or increasing pectin methylation may result in increased cell wall porosity^15^. However, neither cell wall thickness, porosity or mesophyll conductance were measured in these prior studies overexpressing CGR3 or CGR2^26,27^.

We hypothesized that genetically upregulating CGR3 may improve CO_2_ diffusion through the cell wall by decreasing its thickness and increasing its porosity, and further hypothesized this in turn would increase *g*_m_, CO_2_ concentration at Rubisco (*C*_c_) and, leaf CO_2_ uptake rate (*A*). This was tested by engineering *AtCGR3* into tobacco and molecular and physiological phenotyping of the resulting events in controlled environments and in the field as a test of concept.

## Results

### Transgenic tobacco expressing AtCGR3

A construct expressing the *Arabidopsis* pectin methyltransferase CGR3 was designed to test the hypothesis that up-regulating CGR3 will decrease the thickness and increase the porosity of the cell wall to improve mesophyll conductance. This construct contains the *Arabidopsis* ubiquitin 10 promoter and 5’ leader to drive constitutive expression of *AtCGR3.* As antibodies were not available, a C-terminal FLAG epitope tag was included before the *Arabidopsis* heat shock protein 18 terminator (Fig. 1a). The construct was stably transformed into tobacco cv. Samsun, and T2 homozygous plants from three independent single insertion events were characterized in the greenhouse and field. Non-transgenic wildtype (WT) tobacco plants of the genotype transformed and equivalent generation propagated in the same environment were used as controls.

**Fig. 1:**
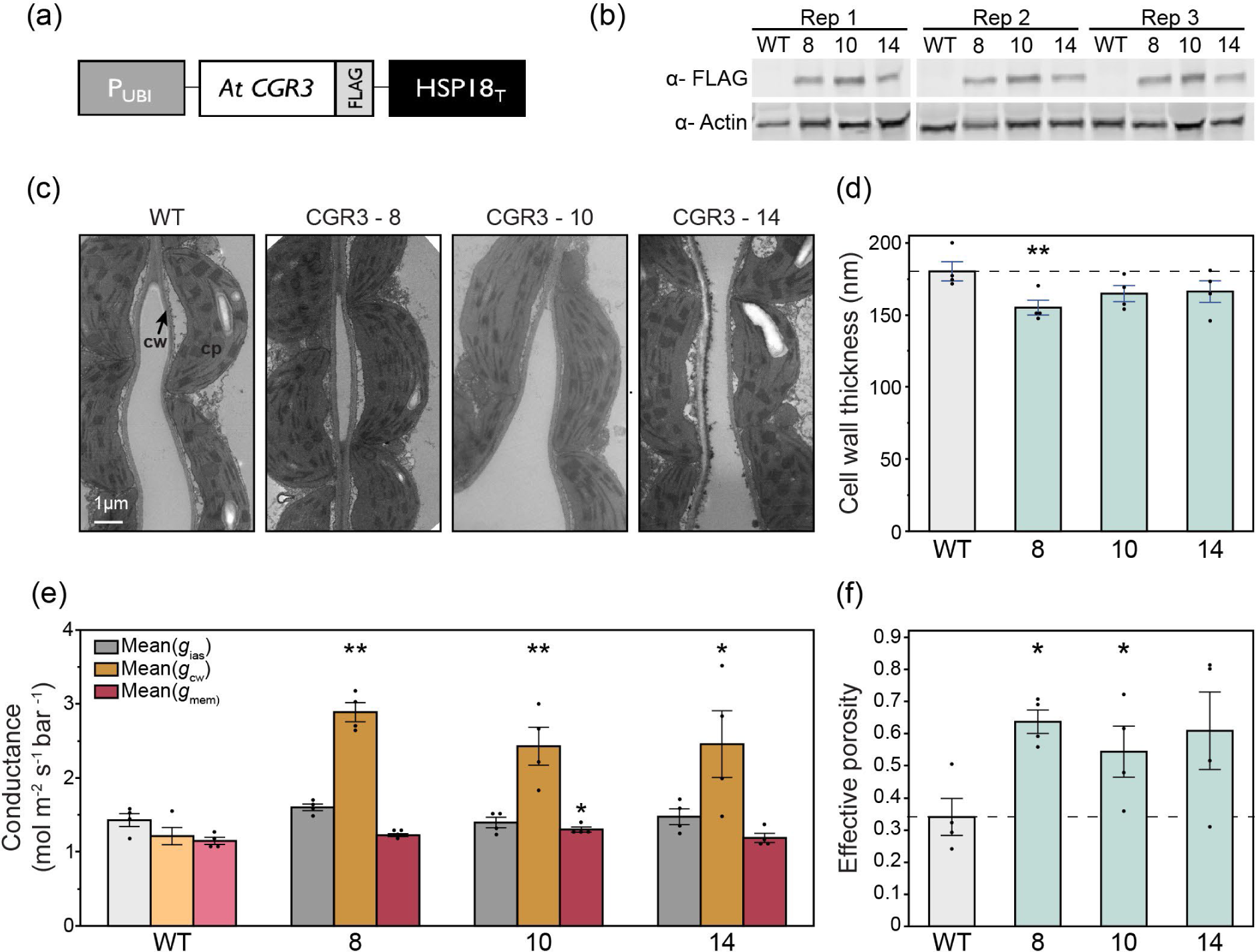
AtCGR3 protein expression in tobacco and its effect on CO_2_ conductance across the cell wall. **(a)** Transgene designed to constitutively express an Arabidopsis pectin methyltransferase CGR3. The transgene was stably transformed into tobacco cv. Samsun. **(b)** Total soluble protein isolated on a leaf area basis from single copy T2 homozygous plants and analyzed by immunoblot. Three transgenic events (8, 10 and 14) and the wildtype (WT) control were probed with anti-FLAG and anti-Actin antibodies. CGR3 protein is ∼28 kDa. Actin was used as a loading control. **(c)** Representative transmission electron micrographs for each event. cw, cell wall; cp, chloroplast. **(d)** Mesophyll wall thickness measured from electron micrographs. **(e)** Estimated CO_2_ conductances across the intercellular airspace (*g_ias_*), cell wall (*g_cw_*), and membranes (*g_mem_*), expressed on a leaf area basis. **(f)** Effective porosity (*p*/*τ*) of the cell wall. Values are shown as the mean ± SEM (n = 4). Asterisks indicate significant differences between WT and the CGR3 transgenic line (**P < 0.05, *P < 0.1); one-way ANOVA, Dunnett’s post hoc test; *g_cw_* significance determined with Welch ANOVA, Games-Howell post hoc test.

qPCR analysis confirmed that all transgenic lines had high levels of *AtCGR3* RNA expression, while no expression was detected in the WT controls (Extended Data Fig. S1). Immunoblotting was then used to ensure *At*CGR3 protein was accumulating in the transgenic tobacco plants. Strong CGR3 protein expression was observed exclusively in the transgenic plants when probed with anti-FLAG (Fig. 1b).

### AtCGR3 expression increases CO_2_ conductance across the cell wall

Mesophyll chloroplast ultrastructure observed by transmission electron microscopy showed no differences between genotypes (Fig. 1c). Mesophyll cell wall thickness (*T*_cw_), was decreased 7-13% in the transgenic plants expressing *AtCGR3* (Fig. 1d).

*g*_m_ includes CO_2_ diffusion across multiple sequential barriers, each of which have an associated conductance *g*. The conductances across the intercellular airspace (*g_ias_*), cell wall (*g_cw_*) and membranes (*g_mem_*) are expected to have the largest effects on *g*_m_ and can be estimated using measured values of *g_m_*, *f_ias_*, *T_cw_*, *T_mes_*, and *S_c_* ^29,30^. Using the corresponding measured values presented in Fig. 1-3, we calculated that plants expressing AtCGR3 had a significantly increased *g_cw_* of 114%, with no significant changes in *g_ias_* or *g_mem_* (Fig. 1e). *g_cw_* is directly influenced by cell wall thickness (Fig. 1d), porosity and tortuosity^15^. Effective porosity (*p*/*τ*) of the cell wall was calculated to have increased by an average 75% compared to the WT control (Fig. 1f).

Representative light micrograph images (Fig. 2a) show differences in leaf and mesophyll thickness (*T*_mes_). CGR3 expression resulted in significant increases in the fraction of intercellular airspace (*f*_ias_) of approximately 12% (Fig. 2b), as well as minor increases in *S*_c_/*S*_m_ in the three independent transgenic events (Fig. 2c). *T*_mes_ was increased in two of the three independent transgenic events (Fig. 2d). No change in leaf mass per unit area (LMA) was observed (Extended Data Table 2).

**Fig. 2:**
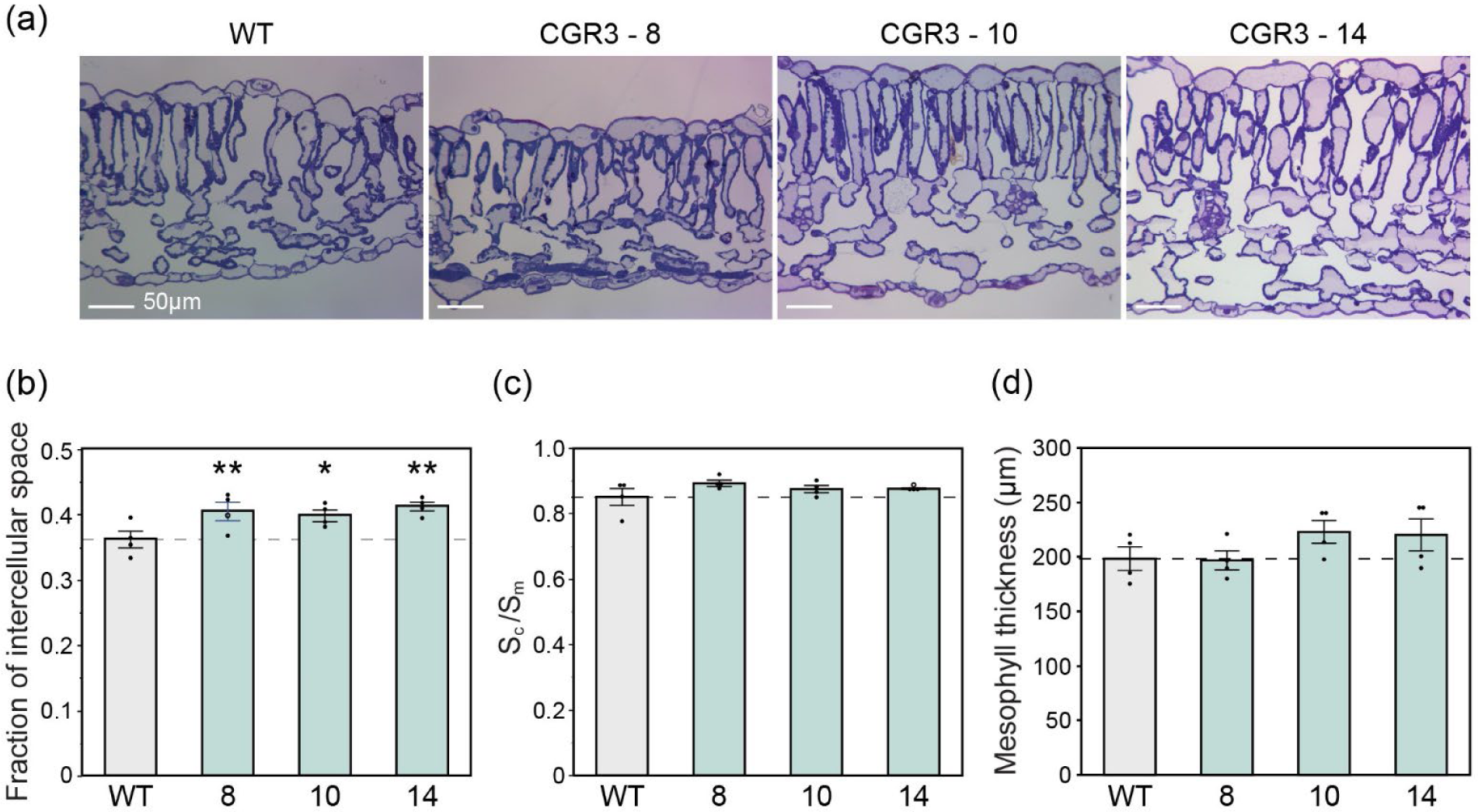
Light micrographs of transverse leaf sections and measured leaf anatomical traits. **(a)** Representative light micrographs. Light micrographs were used to measure **(b)** fraction of intercellular airspace **(c)** ratio of chloroplast surface area exposed to intercellular airspaces (S_c_) to mesophyll surface area exposed to intercellular airspaces (S_m_), and **(d)** mesophyll thickness. Values are shown as the mean ± SEM (n = 4 plants). Asterisks show significant differences between WT and the CGR3 transgenic line (**P < 0.05, *P < 0.1); (b) and (d) one-way ANOVA, Dunnett’s post hoc test; (c) Wilcoxon’s non-parametric test.

**Fig. 3:**
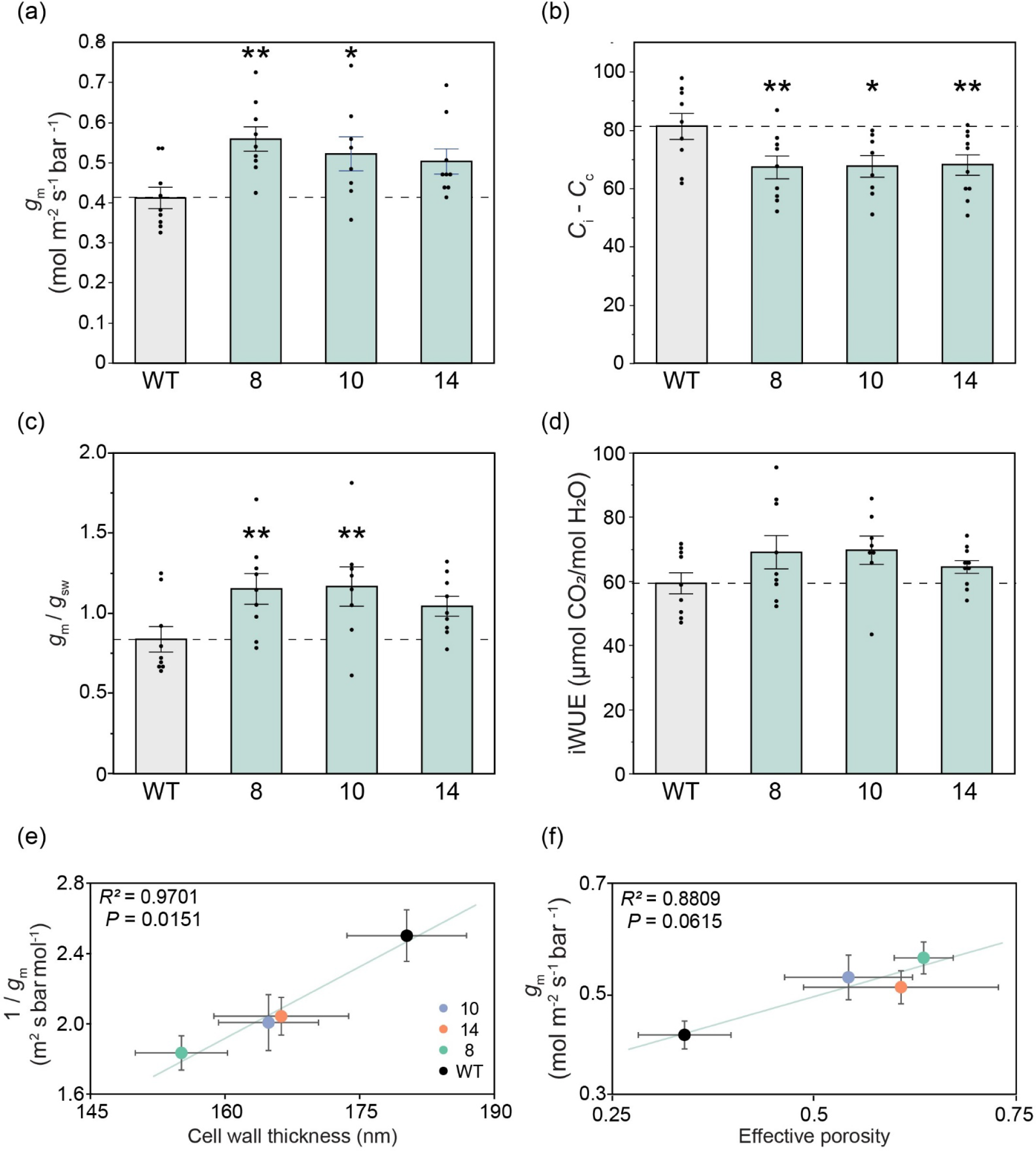
Mesophyll conductance and associated parameters estimated from carbon isotope discrimination (Δ ^13^C) coupled with gas exchange at 2% oxygen in greenhouse grown tobacco. **(a)** Mesophyll conductance (g_m_) calculated from Δ ^13^C, **(b)** the drawdown of CO_2_ into the chloroplast (C_i_ – C_c_), **(c)** the ratio of mesophyll conductance (g_m_) to stomatal conductance (g_sw_) and **(d)** intrinsic water use efficiency (iWUE), the ratio of net CO_2_ assimilation rates (*A*) to stomatal conductance (g_sw_). Measurements were made under the following conditions: light intensity of 1800 µmol m^-2^ s^-1^, leaf temperature of 25 °C, 2% O_2_, and 400 µmol mol^-1^ CO_2._ **(e)** The relationship between 1/g_m_ and mesophyll cell wall thickness and, **(f)** the relationship between g_m_ and effective porosity. The solid lines represent linear regressions from the data points calculated using Pearson’s coefficient of correlation. Values are shown as the mean ± SEM (n = 4).

To explore whether CGR3 expression altered cell wall composition, we measured cell wall pectin, hemicellulose and cellulose content. No primary cell wall component showed any significant differences between CGR3 and WT (Extended Data Table 2). Additionally there was no difference in the ratio of pectin content to the sum of hemicellulose and cellulose content, a value used to indicate cell wall porosity (Extended Data Table 2)^15^.

### Increases in *g*_m_ estimated from Δ ^13^C in transgenic lines grown under controlled growth conditions

Mesophyll conductance was measured to assess whether the anatomical changes described above, including decreased *T*_cw_ and increased *f*_ias,_ affect CO_2_ diffusion. Multiple methods were used to overcome some of the uncertainties associated with estimating *g*_m_. First, carbon isotope discrimination (Δ ^13^C) measurements coupled with gas exchange at 2% oxygen were used to estimate *g*_m_ in greenhouse grown tobacco. Δ ^13^C measurements showed that *g*_m_ was increased in all three events by an average of 28% relative to WT (Fig. 3a). Concomitantly, all three events showed a significantly smaller drawdown of [CO_2_] between the stomatal cavity and chloroplast stroma (*C*_i_ – *C*_c_), averaging a 20% less drawdown than WT and therefore a greater [CO_2_] at Rubisco (Fig. 3b). No changes in stomatal conductance (*g*_sw_) were observed, resulting in significant increases in the ratio of *g*_m_/*g*_sw_ (Fig. 3c). Small increases in CO_2_ assimilation (*A*) were observed, resulting in indicated increases in intrinsic water use (iWUE; Fig. 3d) in all three events, although these were not statistically significant.

Total leaf sugar and starch trended higher in all three transgenic events relative to WT, consistent with increased CO_2_ assimilation (Extended Data Fig. 2). To check for pleotropic effects from increasing *g*_m_, stomatal density and chlorophyll content was measured. All genotypes had similar adaxial and abaxial stomatal densities (Extended Data Fig. 3a-b) and no change in the ratio of abaxial:adaxial stomatal densities was observed (Extended Data Fig . 3c). In addition, leaf chlorophyll content, as measured using a SPAD meter, did not differ between WT and transgenic plants (Extended Data Table 2).

In addition, 1/*g*_m_ was significantly lower in all CGR3 events (Extended Data Table 2) and showed a positive correlation with cell wall thickness (*P* = 0.0151, *R*^2^ = 0.97, Fig. 3e), consistent with previous studies^14,24^. In addition, *g*_m_ was significantly correlated with effective porosity (*P* = 0.0615, *R*^2^ = 0.88, Fig. 3f).

### Increased *g*_m_ in AtCGR3 events confirmed under field conditions using the Variable *J* method

Subsequently a field experiment was conducted to assess whether differences observed in *g*_m_ under greenhouse conditions were reproduced under field conditions. In 2022, a field experiment was carried out with replicated plots of the same three independent transgenic events overexpressing AtCGR3, using a randomized block design (Extended Data Fig. 4a).

Gas exchange measurements were made on the field grown plots to evaluate the physiological effects of decreasing thickness and increasing porosity of the mesophyll cell walls. To test if *g*_m_ was altered, gas exchange measurements were made in parallel with chlorophyll fluorescence measurements. We measured CO_2_ assimilation rates (*A*) as a function of intercellular CO_2_ concentrations (*C*_i_) under saturating light and fit the *A*-*C*_i_ curves using the Variable *J* method to derive *g*_m_^31,32^. This method models the relationship between *A*, the electron transport rate (*J*), and *C*_c_ to estimate *g*_m_ over a range of intercellular [CO_2_] (Fig. 4a). Although this method is generally considered less reliable than those made from Δ ^13^C isotopic measurements^33^, the tunable laser diode system is not field portable. Measurements made at approximately ambient [CO_2_] (400 µmol mol^-1^ CO_2_) showed an average 18% increase in *g*_m_ in the CGR3 transgenic plants that was statistically significant, consistent with the prior greenhouse study (Fig. 4b). Importantly, and consistent with increased [CO_2_] at Rubisco, CO_2_ assimilation rates were significantly increased by an average of 8% in the CGR3 plants relative to the WT controls (Fig. 4c). However, *g*_sw_ was also marginally increased (Fig. 4d), resulting in no significant change in intrinsic water use efficiency in the field. No change in the slope of *A* versus *g*_sw_ was apparent between WT and the transgenic plants (Extended Data Fig. 6a). Both *g*_m_ and *g*_sw_ had a strong positive correlation with CO_2_ assimilation (Extended Data Fig. 6).

**Fig. 4:**
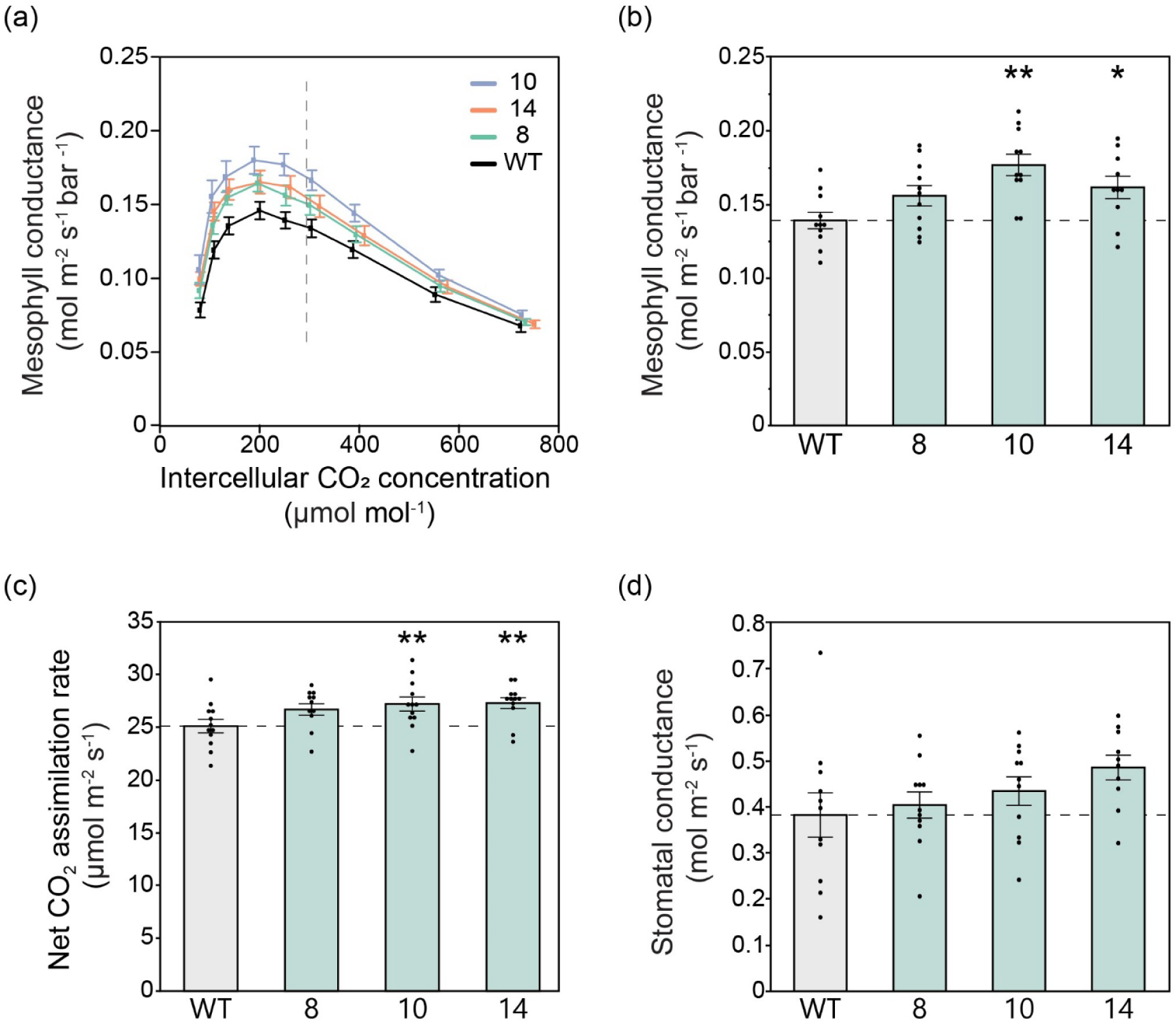
CO_2_ assimilation measured with gas exchange in parallel with chlorophyll fluorescence to estimate mesophyll conductance in field grown tobacco plants. **(a)** Mesophyll conductance (g_m_) as a function of intercellular CO_2_ concentration, estimated using the Variable J method. The vertical dashed line shows the average operating C_i_ (where ambient CO_2_ is 420 µmol mol^-1^). **(b)** Mesophyll conductance measured at 400 µmol mol^-1^ CO_2_ derived from (a). **(c)** Net CO_2_ assimilation rates and **(d)** stomatal conductance to water (g_sw_), each measured at 400 µmol mol^-1^ CO_2._ **a-d** measurements made at light intensity of 1800 µmol m^-2^ s^-1^, leaf temperature of 28 °C, and 60% humidity. Values are shown as the mean ± SEM (n =10-11). Asterisks indicate significant difference between WT and the CGR3 transgenic line (**P < 0.05, *P < 0.1); one-way ANOVA, Dunnett’s post hoc test.

### *V_c,max_* and apparent *V_c,max_* estimates from field gas exchange measurements consistent with increased *g*_m_ in AtCGR3 transgenic plants

The measured *A*-*C*_i_ responses (Fig. 5a) were fit to the Farquhar-von-Caemmerer-Berry model ^34^ to estimate the apparent maximum rate of Rubisco carboxylation (*V_c,max_*) (Fig. 5c). The apparent *V_c,max_* value is determined by the initial phase of the relationship of *A* to intercellular [CO_2_] (*C*_i_), so it is a function of both the actual activity of Rubisco and *g*_m_. To test whether the increase in apparent *V_c,max_* in the transgenic events was the result of increased *g*_m_ or a pleiotropic effect on Rubisco activity, the curves were re-analyzed on a *C*_c_ basis, whose values were derived from the *g*_m_ obtained at each [CO_2_] (Fig. 4a). The initial phase (*V_c,max_*) of the transgenic A-C_c_ curves overlies that of the WT (Fig. 5b), showing that the difference was entirely due to increased *g*_m_ and not Rubisco activity (Fig. 5d).

**Fig 5:**
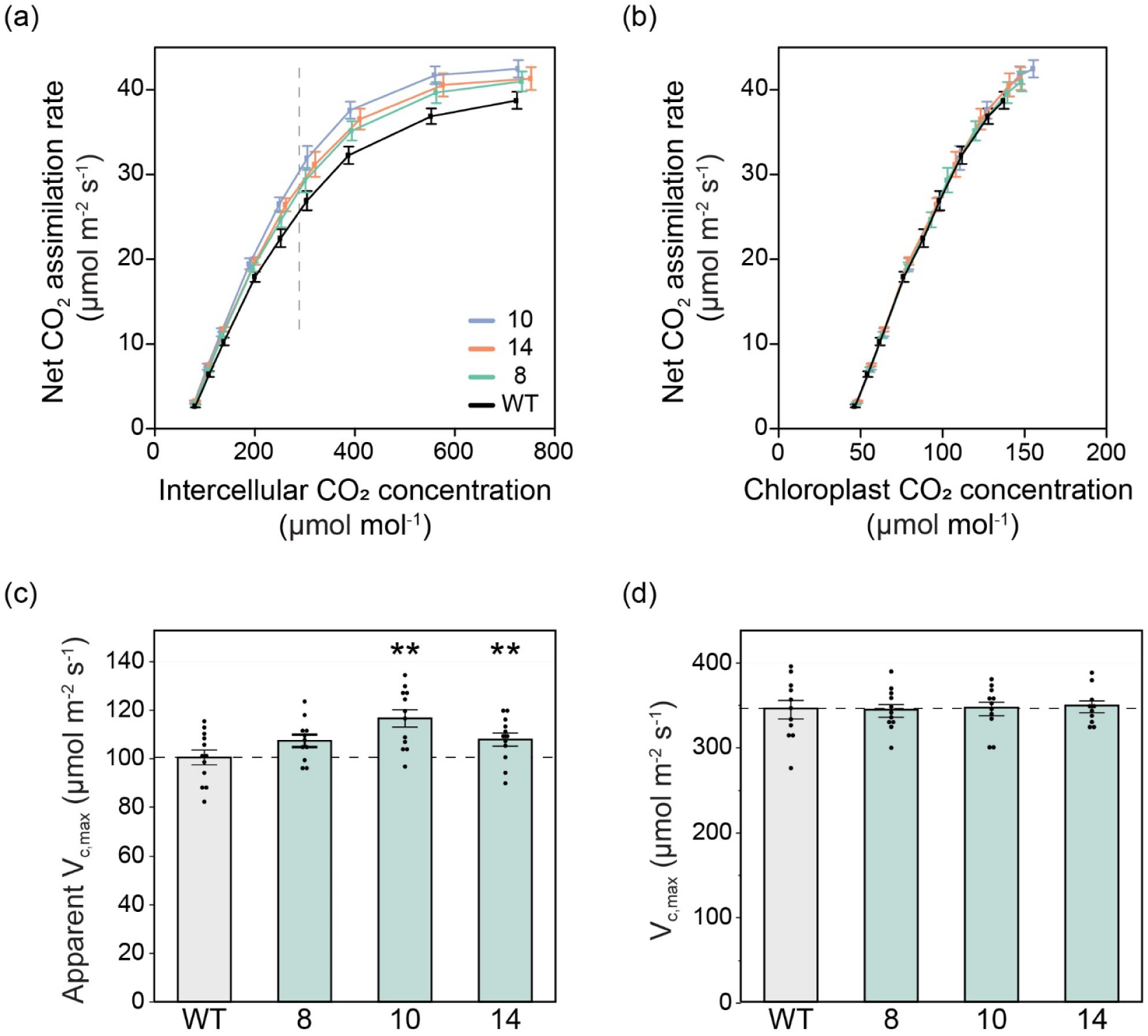
CO_2_ response curves and maximum rates of Rubisco carboxylation based on intercellular [CO_2_] and chloroplast [CO_2_]. **(a)** Response of net CO_2_ assimilation to intercellular [CO_2_] (C_i_). Measurements were made under the following conditions: light intensity of 1800 µmol m^-2^ s^-1^, leaf temperature of 28 °C, and 60% humidity. CO_2_ concentrations varied from 20-1800 µmol mol^-1^ CO_2._ The vertical dashed line is the average operating C_i_ (where ambient CO_2_ is 420 µmol mol^-1^). **(b)** Response of net CO_2_ assimilation to chloroplast [CO_2_] (C_c_). C_c_ estimated from Variable J fits. **(c)** Apparent maximum Rubisco carboxylation rate (V_c,max_) values at 25 °C estimated from response curves in panel A. g_m_ equal to infinity. **(d)** Maximum Rubisco carboxylation rate (V_c,max_) values at 25 °C estimated from response curves in panel B. g_m_ equal to estimated values from Variable J method (Fig. 4a). Values are shown as the mean ± SEM (n =10-11). Asterisks indicate significant differences between WT and the CGR3 transgenic line (**P < 0.05); one-way ANOVA, Dunnett’s post hoc test.

### Biomass maintained in field-grown plants with decreased cell wall thickness and increased porosity

Cell walls function to protect plants from biotic and abiotic stresses as well as provide structural integrity to the plant, facilitate normal growth and play a crucial role in water relations ^35,36^. To assess whether decreased *T*_cw_ had any negative impacts on plant growth and form in the field, we measured a number of plant growth traits (Fig. 6). No significant differences in plant height, leaf area, or total dry biomass were observed between the transgenic lines and control plants (Fig. 6 a-c). In addition, there were no changes in leaf number or biomass of leaves, stems or roots when weighed individually (Extended Data Table 3). We did not observe any differences in structural integrity, lodging, pest or pathogen stress between the AtCGR3 and WT plants; the transgenic lines were essentially indistinguishable from the WT controls (Fig. 6d). These results are consistent with growth measurements made in the greenhouse, with the exception that leaf number was significantly increased in the AtCGR3 lines in the greenhouse (Extended Data Fig. 4).

**Fig. 6:**
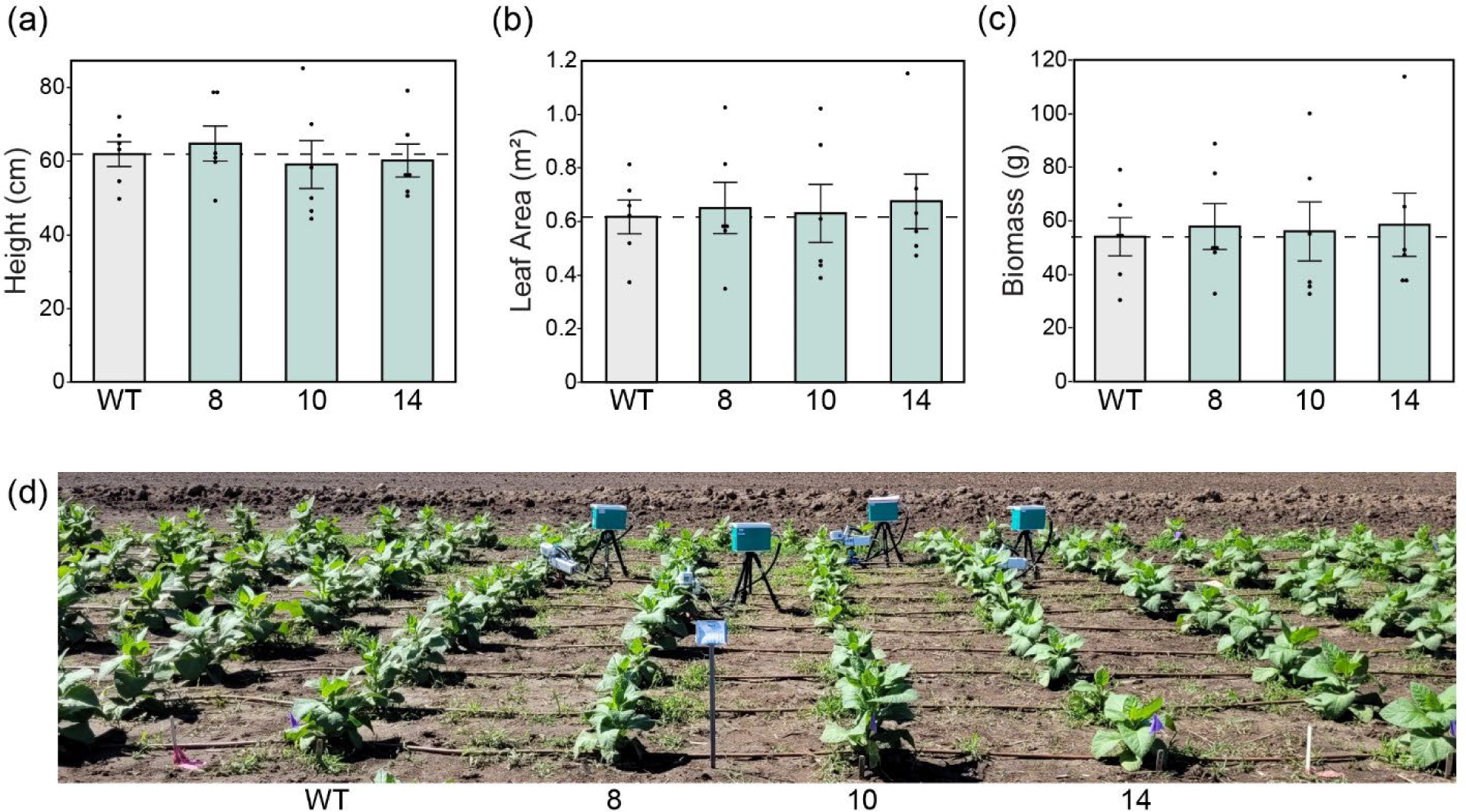
Plant growth traits in field grown tobacco plants. **(a)** Plant height, **(b)** leaf area, and **(c)** biomass (sum of leaf, stem and root dry weights). Values are shown as the mean ± SEM (n = 6 plots). Asterisks indicate significant difference between WT and the CGR3 transgenic line (**P < 0.05, *P < 0.1); (a) and (b) one-way ANOVA, Dunnett’s post hoc test; (c) Wilcoxon’s non-parametric test. **(d)** Tobacco plants growing in the field in Urbana, Illinois summer 2022.

## Discussion

Increasing the diffusive conductance of CO_2_ from the atmosphere to Rubisco has been frequently proposed as an important target for improving CO_2_ assimilation in C_3_ species^6,7,37,38^. Yet, there have been few successes in engineering a change in *g*_m_ into crops. This is at least partly due to an incomplete mechanistic understanding of *g*_m_. While aquaporin channels in the plasma membrane and chloroplast surface area were considered prime targets, manipulations have had no or mixed success for C_3_ species^18,21^. However, observations of variation in both thickness and porosity of the cell wall indicated these as another means to increase mesophyll conductance^15,24^. We identified overexpression of *AtCGR3* as an opportunity to both increase porosity and decrease thickness of the cell wall. Three independent over-expression events in tobacco showed, on average, a 75% increase in porosity and a 10% decrease in thickness of the mesophyll cell walls. This corresponded to a 28% increase in *g*_m_ (estimated using two independent methods) and an 8% increase in leaf CO_2_ assimilation rates, without any pleiotropic effects. This study provides the first report of increased mesophyll conductance via increased porosity and decreased thickness of the cell wall in a dicot species. It also appears one of few demonstrated transgenic increases in mesophyll conductance and leaf photosynthesis of a crop within a replicated field trial^39^. This should serve a proven test-of-concept for further manipulations of the cell wall and application to food crops.

Measured values of *g*_m_ are subject to uncertainty because the trait cannot be determined directly and must be estimated using indirect methods. Thus, it is important to check for consistency across different techniques and growth environments. Here, although absolute values are different, similar relative increases in *g*_m_ were observed in each of the three transgenic events relative to WT, both when estimated in the field from chlorophyll fluorescence and in the greenhouse from isotopic ^13^C measurements (Figs. 3-4). Associated measurements of *A* provide another consistency check. Models predict that increasing *g*_m_ on its own should have a modest positive impact on CO_2_ assimilation rates, as observed here^40^. Further analysis showed that the observed increases in *A* and apparent *V*_c,max_ were entirely explained by the observed increase in *g*_m_ (Fig. 5).

Mesophyll conductance is the net effect of several barriers to CO_2_ diffusion and influenced by several aspects of leaf anatomy. Thus, it is important to identify the main drivers of the observed increases in *g*_m_. To investigate this, measured values of the anatomical traits *f*_ias,_ *T*_mes_ and *S*_c_ were used to calculate CO_2_ conductance across the intercellular airspace, cell wall and membrane (Fig. 1). These results indicated that CGR3 expression increased *g*_m_ via increased CO_2_ conductance across the cell wall, without any change in conductance through the intercellular air space or beyond the wall to Rubisco (Fig. 1). It will remain difficult to verify this until methods are established to directly measure each of these conductances within *g*_m_. There were other changes in the leaf. LMA was slightly decreased, which would be expected with a slightly smaller investment in cell wall, which can represent 70% of leaf dry mass^41^. Cell wall composition analysis showed no differences in total pectin or other cell wall components, consistent with results seen in Arabidopsis, suggesting increases in cell wall effective porosity were due to increased pectin methylation^26^. Glycome profiling of the cell wall could be used to gain more insight on changes in crosslinking within the wall by CGR3 expression and how these alterations affect porosity.

Genetic manipulations can affect multiple traits, making it difficult to identify transgenic manipulations that alter *g*_m_ without pleiotropic changes. The few studies successful in increasing *g*_m_ and *A* have either altered additional traits such as true *V*_c,max_, or are unclear about whether these have been altered, making it difficult to determine if *g*_m_ alone can increase photosynthetic rates^39,42–44^. Here we do not observe any changes to the true *V*_c,max_, i.e. that derived from the response of *A* to *C*_c_ (Fig. 5). Decreasing thickness and increasing porosity of the cell wall could be expected to alter mechanical strength of the plant, plant hydraulics, or stomatal function. In the greenhouse and field, there was no observable evidence of any effect on pest damage or plant form. Stomatal density on the adaxial or abaxial leaf surfaces were unchanged, and there was no significant effect on *g*_sw_ (Table S2; Fig.3). Thus, cell wall thickness and porosity has been successfully modified to increase *g*_m_ and *A* without introducing any apparent unintended pleotropic effects.

Despite the significant increase in *A,* no corresponding change in biomass was found in the field. Here, *CGR3* was fused with the *A. thaliana* UBIQUITIN 10, and so the cell wall changes were likely throughout the plant. It is conceivable that use of the constitutive promoter increased respiration costs in plant tissue other than leaves, constraining any increase in plant growth. Future experiments would ideally use leaf mesophyll specific promoters. Mesophyll specific expression of cell wall properties has been obtained using the Rubisco small subunit 1a (*RBCS1A*) promoter^45^.

A major challenge in increasing crop productivity for food security is availability of water^4^. Agriculture accounts for over 70% of water use, and with rising population and climate change, there is little opportunity to gain further water for agricultural use^46^. The air in the sub-stomatal air spaces of leaves is close to water vapor saturation. This means that while increased stomatal conductance will result in increased water loss, increased mesophyll conductance should not have a direct effect on water vapor loss from the leaf. Among the several different approaches to increasing photosynthesis to support increased crop productivity, increasing *g*_m_ is exceptional in its potential to allow an increase in carbon gain without increased water loss^47^. However, in practice this has not been observed, as *A* and *g*_sw_ are strongly correlated although the mechanistic basis of their interdependence is not well understood^8^. Here, increases in *A* and *g*_m_ in the field-grown plants were balanced by increases in *g*_sw_, and no changes in iWUE were observed (Fig. 4d). It is possible that drought conditions may alter this interdependence, allowing for increased *g*_m_, *A* and iWUE. If true, increased *g*_m_ may be most beneficial for sustaining carbon assimilation of plants grown in water limited environments. Our field plants were subjected to temperatures as high at 35 °C in the field, which would have driven large transpiratory fluxes (Extended Data Fig. S5); however, the field plants were irrigated. A recent study by Pathare *et al.* showed engineered increases in *g*_m_ resulted in increased biomass of rice plants grown under reduced soil water content but not those subjected to ample water ^48^.

Taken together, these results provide a critical proof of concept that increasing *g*_m_ by altering the cell wall is a route for enhancing photosynthetic performance of crops. Specifically, the current study shows modification of thickness and porosity as a viable route to improvements in photosynthesis. Gains in water use efficiency could therefore be achieved by combining this increase in *g*_m_ with decreased *g*_sw_, maintaining the same rate of CO_2_ assimilation while reducing water loss from transpiration. Several approaches have now been identified to allow an engineered or bred decrease in stomatal conductance^49–52^. Stacking increased *g*_m_ with other traits such as increased Rubisco activity also has the potential to further increase photosynthetic efficiency. It will be important to consider that certain engineering strategies will only be viable in specific crop species, such as the one here which only applies to C_3_ dicots. Thus, this work complements previous studies that have modified aquaporins and other aspects of leaf architecture, and extends the engineering “toolbox” available for controlling *g*_m_ to further increase photosynthetic efficiency and growth needed to sustainably increase food production.

## Methods

### Plasmid design and assembly

Vector design and construct assembly followed the genetic syntax of the Phytobrick standard^53^ and Loop assembly by Pollak *et al.* (2019)^54^. All required genetic modules were domesticated for BpiI, BsaI and SapI prior to *de-novo* synthesis through TWIST Bioscience. The nucleotide sequence of *A. thaliana* COTTON GOLGI-RELATED 3 (CGR3; AT5G65810.1) was extracted from The Arabidopsis Information Resource (TAIR10)^55^ and codon optimized for *N. tabacum* (IDT^TM^ Codon Optimization Tool). Original and codon optimized CGR3 sequences can be found in Additional file S1. CGR3 was fused with the *A. thaliana* UBIQUITIN 10 (AT4G05320.2) promoter, including the 5’UTR and first intron, a C-terminal 1x FLAG tag and the *A. thaliana* HEAT SHOCK PROTEIN 18.2 3’UTR and terminator (AT5G59720). The CGR3 cassette was combined with a CaMV35S:BAR selection marker and cloned into the pCsB acceptor backbone (Addgene #136068) prior to electroporation into *A. tumefaciens* C58C1. Complete plasmid sequence was verified using next generation sequencing.

### Plant transformation

*N. tabacum* cv. Samsun leaf-disc transformation was performed following Wang (2015)^56^. The following minor modifications were made to the protocol: fully expanded leaf surfaces were submerged in a sterilization solution for 10 minutes. Sterilized leaf discs were rinsed with sterile de-ionized water and cultured in the pre-culture medium. Explants were further incubated at 24°C, with a 16h light period for 48h. *A. tumefaciens* C58C1 containing the target vector was grown overnight to an OD600 of 1.0-1.5 in YEP. Leaf discs were then co-cultivated on fresh pre-culture medium for 48h. After 48h, leaf discs were transferred to a selection medium and incubated under a 16h light period, followed by sub-culturing every 3-4 weeks. Once the shoots reached around 8-10 cm, they were transferred to rooting medium. All media and solution components are described in Methods S1. Established plants were transferred to soil for acclimatization and maturation in the greenhouse after 3-4 weeks.

### Plant growth – greenhouse conditions

T2 homozygous seeds from 3 independent transgenic events and WT *N. tabacum* cv. ‘Samsun’ seeds from which the transgenics were derived and of the same harvest date were germinated on BM6 growing medium (BM6 All-Purpose, Berger) under greenhouse conditions. Ten days after germination seedlings were transplanted to 9cm x 9cm plastic potting trays. After approximately 2 weeks plantlets were transplanted to 3.8L plastic pots (400C, Hummert International) filled with BM6 growing medium supplemented with 15 cm^3^ of 15-9-12 (N-P-K) granulated slow-release fertilizer (Osmocote Plus, ICL-Growing Solutions). Plants were grown under natural illumination with ∼300 µmol m^-2^ s^-1^ of supplemental light, at 28 °C/12-h days and 22 °C/12 hour nights. Chlorophyll content was measured using a SPAD chlorophyll meter (502, Spectrum Tehcnologies). Leaf mass per area (LMA) was measured from 6 leaf discs each ∼1.3cm^2^ which were dried until constant weight and weights recorded. After approximately 9 weeks of growth tobacco plants were harvested. At harvest leaf number, plant height (equal to stem length) and leaf area (LI-3100C area meter, LI-COR) were measured. Stem and leaves were dried to a constant weight at 60°C and dry weights obtained.

### Transcript and protein expression

Plants were grown under controlled environment greenhouse conditions described above or field conditions described in a following section. Four leaf discs (each ∼1.42 cm^2^) were sampled from the youngest fully expanded leaf of 9 week old plants between 11:30 h and 13:30 h, flash frozen in liquid nitrogen and stored at -80°C until processed. Tissue was disrupted and homogenized (TissueLyser Universal Laboratory Mixer-Mill disruptor 85210, QIAGEN), at 20 Hz for one and a half minutes twice, submerging cassettes in liquid nitrogen before each run. mRNA was extracted using the NucleoSpin RNA Plant Kit (Macherey-Nagel 740949) modified to increase first RA3 buffer wash to 650 µl and an additional 400 µl RA3 buffer wash. RNA quantity and quality was assessed by NanoDrop^TM^ One/OneC (Thermo Fisher Scientific). cDNA was synthesized using the SuperScript^TM^ III First-Strand Synthesis System (Invitrogen) with random hexamers and 8 µl of RNA.

qPCR was conducted in a 20 µl reaction of SsoAdvanced Universal SYBR Green Supermix (Bio-Rad), dilute cDNA, and 500nmol of each primer and annealing temperature of 59 °C on a CFX Connect Real-Time PCR Detection System (Bio-Rad) at 95 °C for two minutes followed by forty cycles of 95 °C for 15 seconds and 59 °C for 30 seconds. Calibrated Normalized Relative Quantities (CNRQ) were calculated using qBase+ software v.3.2 (CellCarta) based on the expression of two reference genes, Actin and GAPDH. Primers were designed according to MIQE guidelines^57^. Primer linear range and efficiency were determined by qPCR on pooled concentrated cDNA from 4 plots serial diluted by 1:3. Primer efficiencies were between 100-103% with a linear range between 0.15 to 333 ng. See Table S1 for primer sequences used in this study.

Total protein was extracted from 4 leaf discs (each ∼1.42 cm^2^) collected and ground as described above. Tissue was mixed with 1X protein buffer (2.5% BME (v/v), 2 % SDS (w/v), 10 % glycerol (v/v), 0.25 M Tris HCl (pH 6.8)), heated to 95°C for 5 minutes and the quantity expressed per unit leaf area. Proteins were separated on 15-well, 4-20 % Mini-PROTEAN® TGX^TM^ Precast Protein Gel (Bio-rad) and transferred onto polyvinylidene difluoride membranes (Bio-Rad) using the TransBlot®Turbo^TM^ Transfer System (Bio-rad) using the fast TGX protocol. Anti-FLAG (Agrisera) and anti-Actin (Agrisera) primary antibodies were incubated at a 1:5000 dilution overnight at 4 °C in phosphate-buffered saline with 1% non-fat dry milk (w/v) and 0.1% Tween-20. Membranes were incubated with IRDye® 800CW Donkey anti-Rabbit IgG secondary antibody (LI-COR) at a 1:10000 dilution at room temperature for 1-2 hours. Immunoblots were imaged at 800nm using the LI-COR Odyssey CLx Infrared Imaging System (LI-COR).

### Microscopy and anatomical measurements

Leaf tissue was collected from the interveinal region of the youngest fully expanded leaves and fixed in 2% glutaraldehyde (Electron Microscopy Sciences, EMS) and 2.5 % paraformaldehyde (Ted Pella Inc). Fixed tissue was stored at 4°C in the dark until being processed for light and transmission electron microscopy (TEM). Samples were post-fixed in 2% osmium tetroxide (EMS) and potassium ferrocyanide (Mallinkckrodt Baker Inc) and then stained overnight in 7% uranyl acetate at 4°C. A graded series of ethanol, ending in 100% ethanol was used to dehydrate the tissue, followed by 100% acetonitrile. The tissue was then infiltrated with 1:1 acetonitrile to Lx112 epoxy mixture (Ladd, Inc), 1:4 and then pure epoxy before hardening at 80°C overnight. For light microscopy blocks were trimmed and sectioned at 0.35 microns, stained with toluidene-blue and basic fuchsin, and viewed with a stereo microscope (BH2, Olympus) coupled with an ocular digital camera (AMT). For electron microscopy blocks were sectioned at 60-90nm for electron microscopy and viewed at 75KV where plate film was scanned in at 3200 dpi (H600, Hitachi).

Light micrographs were used to measure the length of mesophyll cells exposed to intercellular airspace (*L*_mes_), the length of chloroplast exposed to intercellular airspace (*L*_c_) and the width of each section measured (*W*). Mesophyll surface area exposed to intercellular airspace (*S*_mes_) and chloroplast surface area exposed to intercellular (*S*_c_) were calculated using Equations 4 and 6 from Evans *et al.* (1994)^58^.

At least three non-overlapping fields of view were randomly selected to provide technical replicates, which were averaged to provide a single value for each of the four biological replicates for each genotype. Transmission electron micrographs were used to measure mesophyll cell wall thickness. Approximately 6500 nm of cell wall was measured per genotype. 10 non-overlapping fields of view were measured from each of the four biological replicates per genotype. For each image (technical replicate) the area of the cell wall divided by the length was used to calculate cell wall thickness. This accounts for small variations in thickness along the cell wall. Technical replicates were averaged to provide a single value for each biological replicate (4 per genotype). All measurements were made using the freehand area and line selection tools from ImageJ (National Institutes of Health).

### Leaf and Cell wall composition analysis

Fully expanded leaves with midrib excised were flash frozen in aluminum foil packets in liquid nitrogen before storage at -80°C. Three to six grams of tissue were lyophilized to a steady state weight. Lyophilized tissue was ground for 15 minutes at 1200 rpm on a Genogrinder 2010 (SPEX) using two 4 mm stainless steel grinding beads. Total sugars were extracted from 100-200 mg of ground, dried tissue by incubation in 80% ethanol at 80°C for 20 minutes with decantation six times ^59^. Ethanol extracts were treated with activated charcoal to remove compounds such as lactic acid, sugar alcohols, and alcohol-soluble pigments which can interfere in the reaction and lead to overestimations of sugar content. Total sugar as glucose was measured using the sulfuric-phenol microplate assay as described in Kondo *et al.* (2021)^60^. The protocol was modified to change the heat treatment to 90 °C in a water bath for 5 minutes. Sugar extract absorbance at 490 nm was measured in triplicate on a Synergy HI Microplate spectrophotometer (Biotek) against a 5-25 ug glucose standard curve.

After ethanol extraction the remaining pellet was washed with 1:1 chloroform:methanol (v/v), followed by acetone, and dried overnight at 35°C. The pellet was subjected to three rounds of digestion by 500 µL of 120 U/mL α-amylase Bacillus licheniformis (Neogen) in 10 mM, pH 6.5 MOPS buffer at 75 °C for 30 minutes ^59^. The enzyme was deactivated by heating at 99 °C for 10 minutes. After centrifugation at 13,000 g for 10 minutes, 800 µl of supernatant was quantitatively transferred and subjected to two rounds of digestion by 500 µL of 30 U/mL amyloglucosidase Aspergillus niger (Neogen) in 100mM, pH 4.5 acetate buffer at 50 °C for 30 minutes ^59^. Total starch as glucose was measured by D-Glucose GOP-POD microplate assay (nzytech). The pellet from the α-amylase MOPS digestion was decanted, washed with water twice, and acetone once. The acetone was removed using a Speed Vac Concentrator (Thermo Fisher Scientific) to steady state weight resulting in the cell wall alcohol insoluble residue (AIR).

In triplicate, 2-3 mg of AIR was digested in 375 µl 2 M trifluoroacetic acid (TFA) at 121°C for 90 minutes^61^. Supernatant was removed and analyzed for TFA soluble hemicellulose by the sulfuric-phenol microplate assay described previously and for pectin as D-Galacturonic acid per Bethke and Glazebrook (2019)^62^ with minor modifications. The addition of 2 mg/mL prepared m-hydroxydiphenyl reagent was reduced to 10 µl per well and measured absorbance at 525 nm in triplicate on a Synergy HI Microplate spectrophotometer (Biotek) against a 6.25 to 200 nmol D-(+)-galacturonic acid monohydrate (AAJ6628214, Thermo Fisher Scientific) standard curve.

Cellulose and non-soluble hemicellulose (primarily glycan) was digested with sulfuric acid as described in Foster, Martin, and Pauly (2010)^61^ and quantified as glucose by the sulfuric-phenol microplate assay described previously against a 2-12 µg glucose standard curve. Detailed protocol available at protocols.io. https://dx.doi.org/10.17504/protocols.io.3byl4q6jzvo5/v1.

### Stomatal density

Adaxial and abaxial stomatal impressions of approximately 2 cm^2^ were made on the youngest fully expanded leaf of greenhouse grown plants as described previously^63^. Six plants per genotype were sampled. Four images were obtained per impression using the Axio Imager A1 microscope (Zeiss) equipped with the Zeiss AxioCam HrC digital camera, AxioVision software version 4.9.1.0 (Zeiss) and a 20x/0,5 objective (EC Plan-Neofluar420350-9900). All whole stomata and partial stomata on the left and top borders of the image were counted using Cell Counter Plugin (https://imagej.net/ij/plugins/cell-counter.html) in ImageJ ^64^ and used to calculate stomatal density.

### Estimating mesophyll conductance using carbon isotope discrimination coupled with leaf gas exchange

The LI-COR 6800 gas exchange system (LI-COR Environmental) was coupled to a tunable-diode laser absorption spectroscope (TDLAS model TGA 200A; Campbell Scientific) to measure online carbon isotope discrimination ^65,66^. The TDL was connected to the LI-6800 reference and sample air streams using the ports on the back of the sensor head. N_2_ and O_2_ were mixed using mass flow controllers (OMEGA Engineering Inc.) and spilt into multiple lines to use as CO_2_ free air. One line was used to zero the TDL throughout the measurements. Two lines supplied the inlets of two gas exchange systems to make measurements at 2% O_2._ The final line was diluted with a 10% CO_2_ gas cylinder to produce three different CO_2_ concentrations (60, 300 and ∼1000 ppm CO_2_) of the same isotopic signature and used to calibrate the ^13^CO_2_ signal.

The measurements cycled through nine gas streams in the following sequence: calibration zero, calibration points 60, 300 and 1000 ppm CO_2_, NOAA calibration of δ^13^C composition (NOAA Global Monitoring Laboratory), LI-COR 6800 #1 reference and leaf chamber air streams, and LI-COR 6800 #2 reference and leaf chamber air streams. Each step had a duration of 20 s and measurements were averaged over the last 10 s to produce a single data point.

Gas exchange measurements were made under the following conditions: light intensity of 1800 µmol m^-2^ s^-1^, leaf temperature of 25°C, leaf vapor pressure deficit of 1.3 kPa, 2% O_2_, and 400 µmol mol^-1^ CO_2._ Two percent oxygen was used to minimize photorespiration. Once steady-state CO_2_ assimilation and stomatal conductance were reached the gas-exchange system was set to auto-log at 180 s intervals over the course of 30 min. After the program was completed, the light was turned off and dark respiration rate was measured on plants after >30 minutes in the dark.

The combined gas exchange and TDLAS data were processed and analyzed using PhotoGEA, an R package for photosynthetic gas exchange analysis ^67^. This process generally followed the steps described in the “Analyzing Mesophyll Conductance Data” article included with PhotoGEA, which is also available online at the PhotoGEA documentation website: https://eloch216.github.io/PhotoGEA/.

Within each TDL cycle, correction factors derived from the five calibration tanks were used to obtain calibrated dry-air [^12^CO_2_] and [^13^CO_2_] in the air streams entering and exiting each LI-COR leaf chamber. The isotopic composition (δ^13^C) of each air stream was calculated using Equation 4 from Ubierna *et al*. (2018)^68^. Timestamps and TDL valve numbers were then used to pair each TDL measurement with its corresponding gas exchange log entry, enabling the calculation of the observed photosynthetic ^13^CO_2_ discrimination (Δ^13^C) and the ternary gas correction factor (*t*) using Equations 5 and 9 from Ubierna *et al*. (2018) ^68^. The CO_2_ compensation point in the absence of day respiration (Γ*) was calculated from [O_2_] and leaf-temperature-dependent O_2_ and CO_2_ solubilities assuming a Rubisco specificity of 97.3 M M^-1^ ^69^. Finally, mesophyll conductance to CO_2_ diffusion (*g_mc_*) was calculated using Equations 13 and 22 from Busch *et al*. (2020) ^70^, which assume that mitochondrial respiration is isotopically disconnected from the Calvin-Benson-Bassham cycle. The effective isotopic fractionation due to day respiration (*e\**) was calculated using Equation 19 from Busch *et al*. (2020) ^70^ rather than Equation 20, because values of 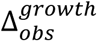 were not available; however, this should have minimal impact due to the low [O_2_] used for these measurements.

### Plant growth – field conditions

Seeds from homozygous T2 single insertion events (CGR3-8, CGR3-10 and CGR3-14) and WT seed from the same harvest date were sown in the greenhouse on May 16^th^ 2022 (DOY 136). After 10 days seedlings were transplanted to floating trays as described in Kromdijk *et al.* (2016)^71^. Plantlets were transplanted to the University of Illinois Energy farm field site (40.11°N, 88.21°W, Urbana, IL, USA) on June 10^th^ 2022 (DOY 161). The field was prepared one week prior to transplant as described previously^71^.

The field experiment used a randomized block design with six blocks. Each block consisted of 4 rows of 10 plants per genotype in a north-south (N-S) orientation, with plants spaced 61 cm apart (Extended Data Fig. 5a). Each block contained one WT row. In addition, one border row of WT plants surrounded the perimeter of the 6 experimental blocks. Plants were irrigated as needed using parallel drip irrigation lines (DL077, The Drip Store). Weather data were measured with a digital sensor mounted 10 m above ground level at the same field site (ClimaVUE50, Campbell Scientific, Extended Data Fig. 5b-c).

Plants were harvested July 21^st^ 2022 (DOY 202). At harvest leaf number, plant height (equal to stem length) and leaf area (LI-3100C area meter, LI-COR) were measured. Harvested material was partitioned into leaf, stem and roots for 5 randomly selected plants per row. These were dried to a constant weight at 60°C in custom built dying ovens and dry weights obtained.

### Leaf gas exchange in the field

Photosynthetic gas exchange measurements were performed on the youngest fully expanded leaves on July 9^th^ - 10^th^ 2022 (DOY 190-191). CO_2_ response curves (A-C_i_) were measured using a LI6800 infrared gas exchange system with integrated leaf chamber fluorometer (LI-COR). Leaves were clamped into a 6 cm^2^ gas exchange cuvette and acclimated to the following conditions: light intensity of 1800 µmol m^-2^ s^-1^, leaf temperature of 28 °C, CO_2_ reference concentration of 400 µmol mol^-1^ and 60% humidity. CO_2_ responses were initiated when rates of CO_2_ assimilation and stomatal conductance stabilized to a steady state (∼20 min). Response curve were measured with the following sequence of reference [CO_2_]: 400, 300, 200, 150, 75, 50, 20, 400, 400, 500, 600, 800, 1000, 1200, 1500, and 1800 µmol mol^−1^. Measurements were logged 3 to 5 minutes after each new [CO_2_] step.

Apparent maximum Rubisco carboxylation rates (*V_c,max_*) at 25 °C were estimated using the *fit_c3_aci* function from the PhotoGEA R package ^67^, which fits measured CO_2_ response curves with the Farquhar-von-Caemmerer-Berry (FvCB) model, including limitations from triose phosphate utilization (TPU) ^34^. Temperature scaling of key parameters (*K_C_*, *K_O_*, Γ*, *V_c,max_*, *J*, and *R_d_*) was modeled using Arrhenius factors ^72^ and mesophyll conductance was set to infinity (equivalent to setting *C_c_* = *C_i_*). During the fits, an optimization algorithm is used to choose values of the four unknown FvCB model parameters (*V_c,max_*, *J*, and *R_d_* at 25 °C and the maximum rate of TPU, *T_p_*) that produce the best agreement between the modeled and measured CO_2_ assimilation rates.

### Estimating mesophyll conductance using Variable *J*

*C_c_*, *g_mc_*, and the true *V_c,max_* were estimated from gas exchange measurements made in parallel with chlorophyll fluorescence measurements using the “Variable *J*” fitting method as implemented in the *fit_c3_variable_j* function from the PhotoGEA R package ^67^. In this method, net CO_2_ assimilation (*A_n_*) is modeled by (1) calculating *g_mc_* and *C_c_* from the incident photosynthetically active photon flux density (*Q_in_*), the measured operating efficiency of photosystem II (φ_PSII_), and the measured *A_n_*, and then (2) using the calculated *C_c_* as an input to the FvCB model ^31,32^. There are five unknowns in the equations used to model *A_n_*: τ (a proportionality factor that relates *Q_in_* and φ_PSII_ to the fluorescence-based estimate of the RuBP regeneration rate) and the four FvCB model parameters (*V_c,max_*, *J*, and *R_d_* at 25 °C and *T_p_*). During the fits, an optimization algorithm is used to choose values of these unknowns that produce the best agreement between the measured and modeled *A_n_*. Once these parameter values have been found, values of *C_c_* and *g_mc_* are also immediately known.

### Estimation of effective porosity

The cell wall effective porosity (*p* / *τ*) can be determined from the cell wall conductance to CO_2_ diffusion (*g_cw_*) provided the cell wall thickness *T_cw_* is known ^73^. In turn, *g_cw_* can be estimated from *g_mc_* by accounting for the effect of other known barriers to CO_2_ diffusion (specifically, the intercellular airspace, the plasma membrane, and the chloroplast envelope) ^30,73^. Here we use this approach to calculate *p* / *τ* from measured values of *g_mc_*, *f_ias_*, *T_cw_*, *T_mes_*, and *S_c_*. Overall, our method is similar to the one used in Ellsworth *et al*. (2018) ^73^, but differs by including the conductance across the intercellular airspace and a membrane conductance enhancement factor as in Xiong (2023)^30^. For details of the calculations, see Methods S2.

### Statistical analysis

Normality of the data was tested with Shapiro-Wilk’s test, and homoscedasticity with Brown-Forsythe’s test. If criteria for normal distributions and equal variance was met one-way ANOVA followed by Dunnett’s *post hoc* test for transgenic mean comparison against the WT control was performed. Data were considered significant at *P* < 0.05 and marginally significant at *P* < 0.1. If criteria for normality was violated, Wilcoxon’s non-parametric test was applied. If criteria for equal variance was violated Welch’s ANOVA followed by Games-Howell post hoc test was applied. Analysis of field growth traits (Fig. 6) was performed using a randomized block design with 6 blocks. Tests used are indicated in the figure or table legend. Correlations between 1/*g*_m_ and T_cw_, and *g*_m_ and effective porosity were evaluated using Pearson’s correlation coefficient. Jmp pro version 17.0.0 software was used for all statistical analyses.

## Acknowledgements

We thank Mike Root for the transformation and tissue culture of tobacco. We thank David Drag, Ben Harbaugh, Ben Thompson, Ron Edquilang and Andrew Wszalek for help with the maintenance of greenhouse plants, field experiment and field harvest. We also thank Noga Adar, Maddie Burke and Amanda Eggers for technical lab support. The microscopy work was carried out in part in the Materials Research Laboratory Central Research Facilities, University of Illinois. This work supported by the Bill & Melinda Gates Foundation, Foundation for Food and Agriculture Research and the UK Foreign, Commonwealth & Development Office under grant no. OPP11722157, and from Bill & Melinda Gates Agricultural Innovations grant investment ID 57248.

## Author Contributions

CES and SPL designed the experiments. BEH assembled the construct and supervised generation of the transgenic tobacco lines. SSS set up the TDL and maintained the carbon isotope discrimination equipment. LD measured gene expression, cell wall composition and stomatal density. EBL calculated effective porosity, *g_ias_*, *g_cw_*, and *g_mem_*, and developed the PhotoGEA data-processing R package used to analyze all gas exchange and carbon isotope discrimination data. CES participated in all experiments and analyzed the data. CES and SPL wrote the manuscript with contributions from all authors.

## Extended Data and Supporting Information

Extended Data Fig. 1| Gene expression in field grown CGR3 and WT lines.

Extended Data Fig. 2| Sugar and starch content of greenhouse grown tobacco plants.

Extended Data Fig. 3| Stomatal density of greenhouse grown tobacco plants.

Extended Data Fig. 4| Plant growth traits in greenhouse grown tobacco plants.

Extended Data Fig. 5| Tobacco field experimental design and weather conditions.

Extended Data Fig. 6| Correlation of CO_2_ assimilation to stomatal and mesophyll conductance.

Extended Data Table 1: qPCR primer information

Extended Data Table 2: Summary of leaf gas exchange combined with carbon isotope discrimination, cell wall composition, leaf mass per area (LMA) and chlorophyll content (SPAD value) of greenhouse grown plants.

Extended Data Table 3: Summary of harvest measurements from field-grown plants.

Supplementary Note 1. Plant transformation culture media and solutions components.

Supplementary Note 2. Details for estimation of effective porosity

Supplementary Dataset 1. Codon optimized sequence of AtCGR3

