## Supplemental Figures 1-6, Tables 1-3, Methods 1-2 and Dataset 1 for "Greater leaf photosynthesis in the field by increasing mesophyll conductance via modified cell wall porosity and thickness in tobacco"

### Extended Data

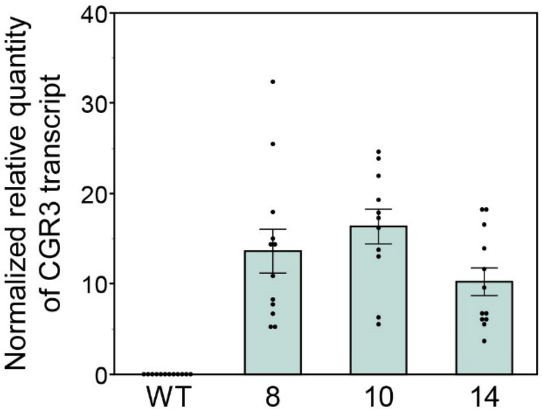


**Extended Data Fig. 1| Gene expression in field grown CGR3 and WT lines.**

qPCR analysis of AtCGR3 gene expression in three independent transgenic events (8, 10, and 14). Values are shown as the mean $\pm$ SEM (n = 12).

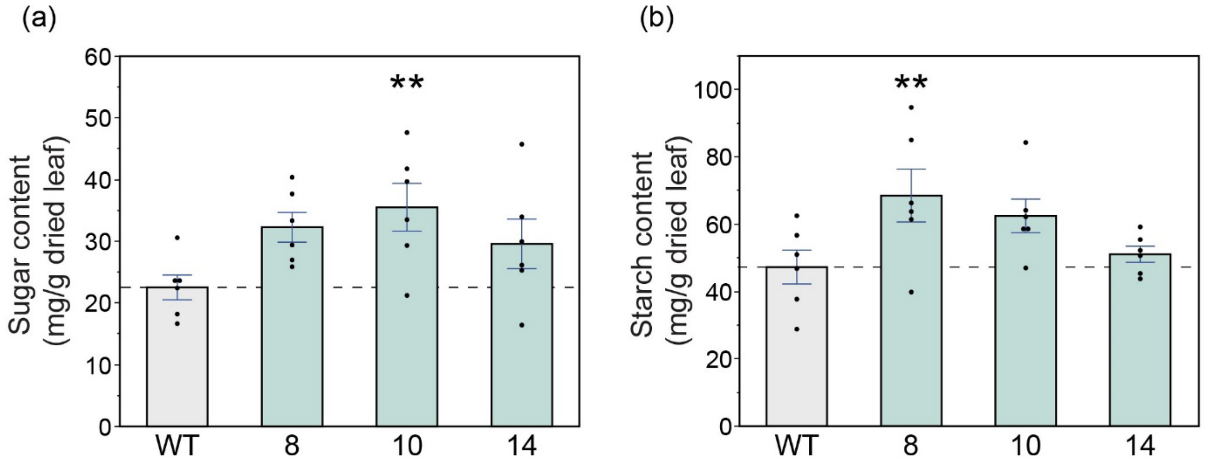


**Extended Data Fig. 2| Sugar and starch content of greenhouse grown tobacco plants.**

Total **(a)** sugar and **(b)** starch content as glucose measured in dried leaf tissue. Values are shown as the mean $\pm$ SEM (n = 6). Asterisks indicate significant differences between WT and the CGR3 transgenic line (**$P < 0.05$); one-way ANOVA, Dunnett's post hoc test.

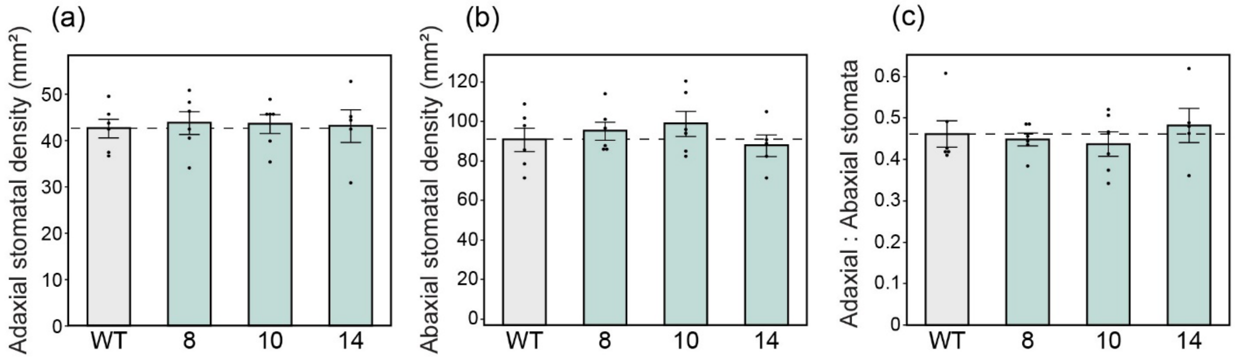


**Extended Data Fig. 3| Stomatal density of greenhouse grown tobacco plants.**

**(a)** Stomatal density on adaxial leaf surface, **(b)** stomatal density on abaxial leaf surface, and **(c)** adaxial to abaxial stomatal density ratio. Values are shown as the mean $\pm$ SEM (n =6). No significant differences, one-way ANOVA, Dunnett's post hoc test.

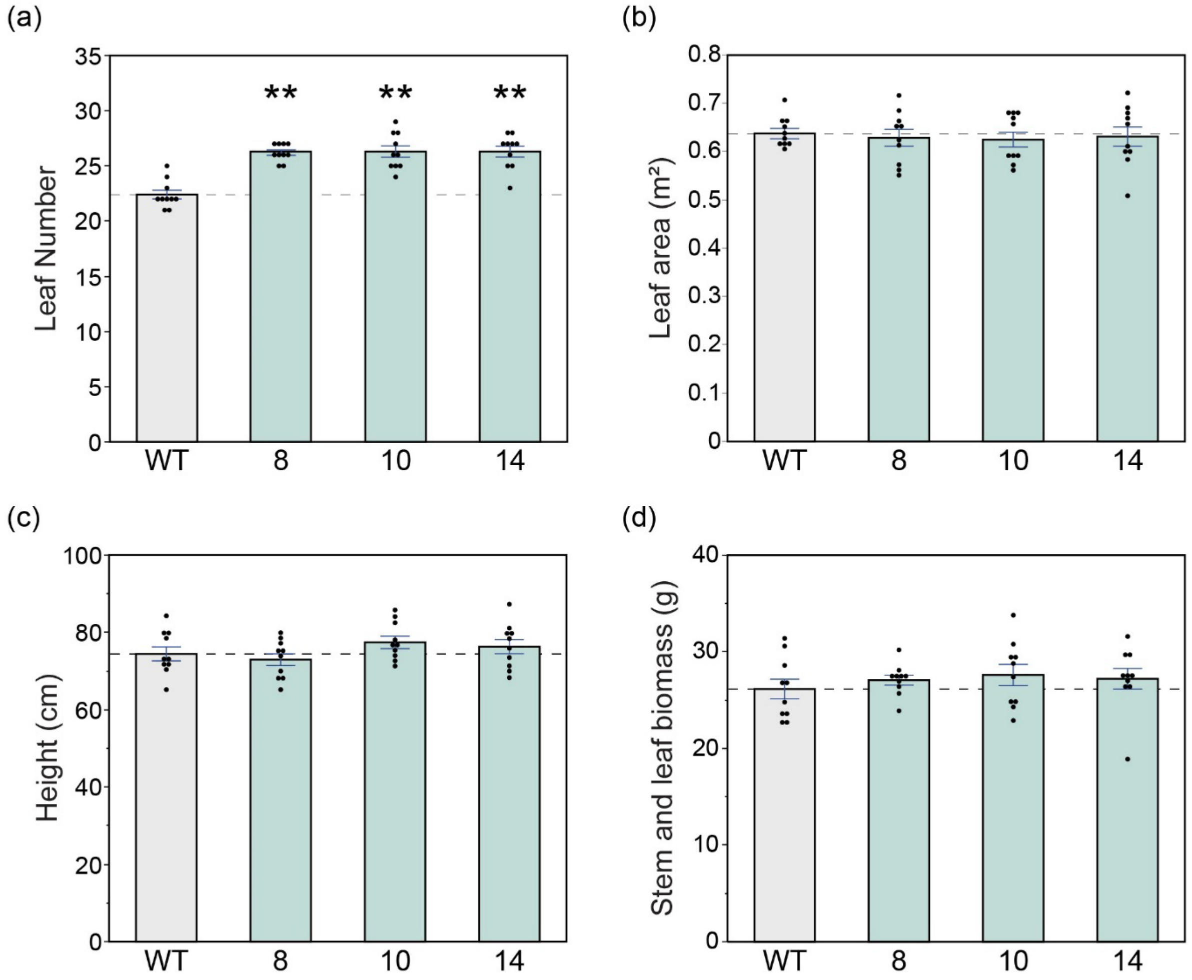


**Extended Data Fig. 4| Plant growth traits in greenhouse grown tobacco plants.**

**(a)** Leaf number, **(b)** leaf area, **(c)** plant height and **(d)** biomass (sum of leaf and stem dry weights). Values are shown as the mean ± SEM (n = 10). Asterisks indicate significant differences between WT and the CGR3 transgenic line (**P < 0.05); (a) Wilcoxon's non-parametric test; (b-d) one-way ANOVA, Dunnett's post hoc test.

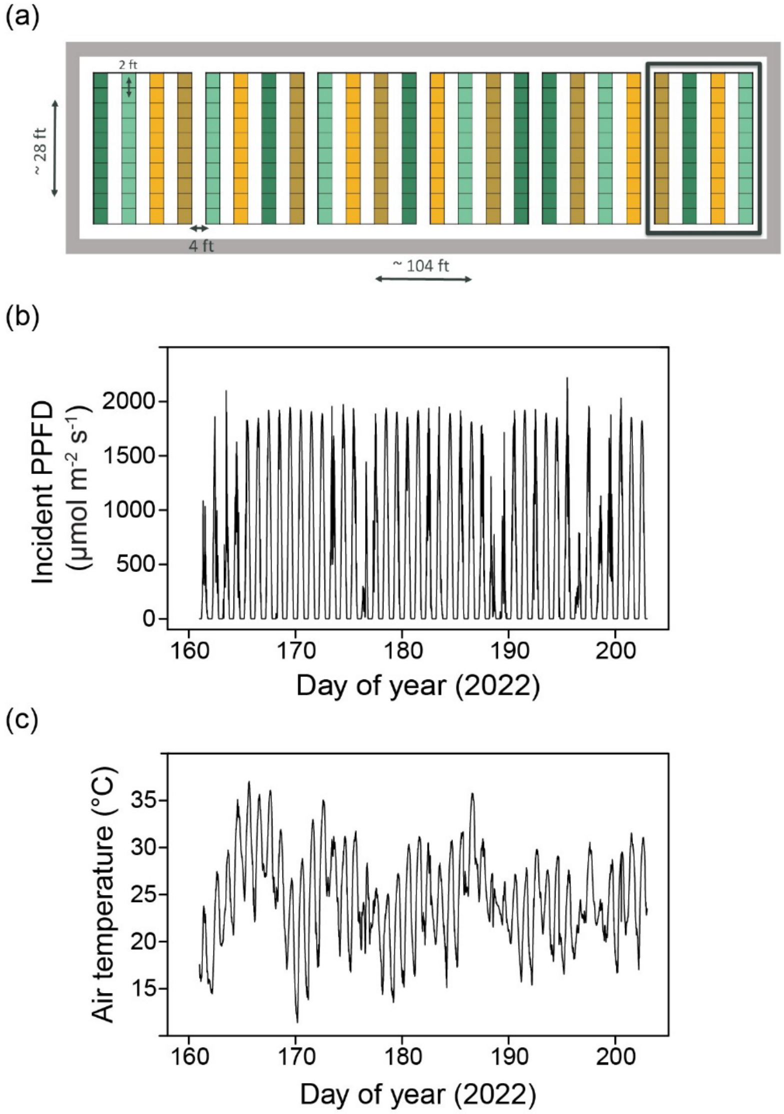


**Extended Data Fig. 5| Tobacco field experimental design and weather conditions.**

**(a)** Schematic of field experimental set up. Genotypes were randomly assigned to one of four possible row positions in each of the six blocks. **(b)** Light intensity from June 10$^{th}$ (DOY 161 - date of tobacco transplant into the field) until July 21$^{st}$, 2022 (DOY 202 - date of tobacco plant harvest). **(c)** Air temperature from June 10$^{th}$ until July 21$^{st}$, 2022 (DOY 161 to 202). Data are shown as averages over the preceding hour.

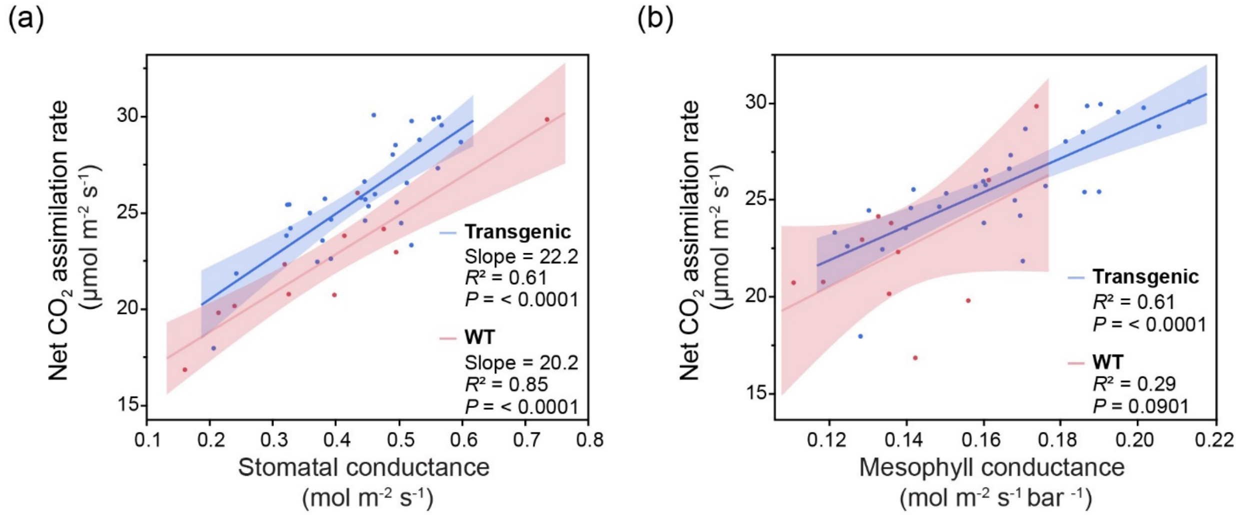


**Extended Data Fig. 6| Correlation of $CO_2$ assimilation to stomatal and mesophyll conductance in field-grown plants.**

The relationship between $CO_2$ assimilation and **(a)** stomatal conductance, or **(b)** mesophyll conductance. Each point corresponds to the values measured from one plant. Measurements from transgenic (blue) and WT (red) plants are shown. The solid lines represent linear regressions from the data points calculated using Pearson's coefficient of correlation. The blue and red areas are 95% confidence intervals of the regression lines.

### Extended Data Table 1 qPCR primer information

| Primer name | Sequence | Amplicon length |
|---|---|---|
| CGR3_F | GTGATCGTCTCCGATGCACT | 80 |
| CGR3_R | CTTGCAACACGAGCCAGTTC | |
| GAPDH_F | CAA CCC CTA ACG TCT CGG TC | 92 |
| GAPDH_R | AGCAGCCTCCCTAAATGCAG | |
| Actin_F | CCT CAC AGA AGC TCC TCT TAA TC | 137 |
| Actin_R | ACA GCC TGA ATG GCG ATA TAC | |

**Extended Data Table 2. Summary of leaf gas exchange combined with carbon isotope discrimination measurements not presented in main data, cell wall composition, leaf mass per area (LMA) and chlorophyll content (SPAD value) of greenhouse grown plants.** Values are shown as the mean ± SEM. Values significantly different from WT at $P < 0.05$ are in bold. One-way ANOVA, Dunnett's post hoc test.

| Parameter | WT | CGR3 – 8 | CGR3 – 10 | CGR3 – 14 | Sample number ($N$) |
|---|---|---|---|---|---|
| $A$ ($\mu mol\ m^{-2}\ s^{-1}$) | 31.37 ± 0.98 | **35.64 ± 1.13** | 33.45 ± 1.20 | 32.93 ± 0.81 | 8-10 |
| $1/g_m$ ($mol\ m^{-2}\ s^{-1}$ bar$^{-1}$) | 2.50 ± 0.15 | **1.83 ± 0.10** | **2.00 ± 0.16** | **2.04 ± 0.11** | 8-10 |
| $g_{sw}$ ($mol\ m^{-2}\ s^{-1}$) | 0.537 ±0.023 | 0.533 ± 0.032 | 0.489 ± 0.023 | 0.515 ± 0.020 | 8-10 |
| Pectin (mg/g AIR) | 86.90 ± 7.42 | 82.92 ± 2.15 | 68.94 ± 5.83 | 69.02 ± 12.4 | 6 |
| Hemicellulose (mg/g AIR) | 157.13 ± 6.52 | 144.18 ± 10.3 | 137.11 ± 14.8 | 159.29 ± 5.05 | 6 |
| Cellulose (mg/g AIR) | 54.22 ± 2.19 | 51.36 ± 5.42 | 47.59 ± 3.36 | 64.94 ± 4.37 | 6 |
| Pectin / (Hemicellulose + Cellulose) | 0.412 ± 0.035 | 0.429 ± 0.025 | 0.302 ± 0.050 | 0.308 ± 0.055 | 6 |
| LMA (g/m$^2$) | 30.27 ± 2.21 | 27.13 ± 1.81 | 28.40 ± 2.02 | 28.89 ± 1.06 | 5-6 |
| SPAD | 55.30 ± 2.47 | 51.52 ± 2.28 | 56.57 ± 2.35 | 52.28 ± 0.756 | 5-6 |

**Extended Data Table 3. Summary of harvest measurements from field-grown plants.** Values are shown as the mean ± SEM. Values are not significantly different from WT at $P < 0.05$.

| Parameter | WT | CGR3 – 8 | CGR3 – 10 | CGR3 – 14 | Sample number (*N*) |
|---|---|---|---|---|---|
| Leaf number | 30.9 ± 3.32 | 31.1 ± 2.79 | 32.8 ± 4.55 | 31.3 ± 4.30 | 6 plots |
| Leaf weight (g) | 29.7 ± 3.21 | 30.9 ± 3.37 | 31.5 ± 4.53 | 33.5 ± 5.55 | 6 plots |
| Stem weight (g) | 13.8 ± 2.03 | 15.0 ± 2.46 | 14.3 ± 3.23 | 14.6 ± 3.17 | 6 plots |
| Root weight (g) | 10.6 ± 2.22 | 11.9 ± 2.89 | 10.2 ± 3.44 | 10.4 ± 3.16 | 6 plots |
| Above ground biomass (g) | 43.4 ± 5.14 | 45.9 ± 5.78 | 45.8 ± 7.73 | 48.0 ± 8.68 | 6 plots |

**Supplementary Information**

**Supplementary Note 1. Plant transformation culture media and solution components.**

Sterilization solution
10% (v/v) Bleach (Sodium hypochlorite)
0.02% Tween 20® (v/v) (Sigma-Aldrich P1379)
1 ml/L Plant Preservative Mixture (PPM™, Plant Cell Technology Inc.)

Pre-culture medium
Murashige and Skoog complete medium with vitamins (Phytotech labs M519)
3% D-Sucrose (Phytotech labs S829)
2 mg/L Kinetin (Phytotech labs K750)
1.0 mg/L Indole-3-Acetic Acid (IAA, Phytotech labs I885)
0.2% Gelzan™ (Phytotech labs G3251)
pH adjusted to 5.6 with 1N KOH and made to required volume

Selection medium
Murashige and Skoog complete medium with vitamins (Phytotech labs M519)
3% D-Sucrose (Phytotech labs S829)
2 mg/L Kinetin (Phytotech labs K750)
1.0 mg/L Indole-3-Acetic Acid (IAA, Phytotech labs I885)
25mg/L Vancomycin (Phytotech labs V8370)
400mg/L Timentin (Phytotech labs T869)
2.0 mg/L Phosphinothricin (Gold Bio P-165-250)
0.2% Gelzan™ (Phytotech labs G3251)

Rooting media
Murashige and Skoog complete medium with vitamins (Phytotech labs M519)
3% D-Sucrose (Phytotech labs S829)
400mg/L Timentin (Phytotech labs T869)
2.0 mg/L Phosphinothricin (Gold Bio P-165-250)
0.2% Gelzan™ (Phytotech labs G3251)

**Supplementary Note 2. Details for estimation of effective porosity**

The full path taken by $CO_2$ from the ambient air to Rubisco includes eight sequential barriers, each of which has an associated conductance $g$: the boundary layer ($g_{bl}$), stomata ($g_{st}$), intercellular airspace ($g_{ias}$), cell wall ($g_{cw}$), plasma membrane ($g_{pm}$), cytosol ($g_{cyt}$), chloroplast limiting membranes ($g_{clm}$), and chloroplast stroma ($g_{stroma}$) (Nobel 2009). By convention, the final six components are often grouped together into a combined mesophyll conductance ($g_m$). Within the mesophyll components, other groupings can also be formed, such as the liquid phase conductance $g_{liq}$ (consisting of $g_{cw}$, $g_{cyt}$, and $g_{stroma}$) and the membrane conductance $g_{mem}$ (consisting of $g_{pm}$ and $g_{clm}$) (Evans and von Caemmerer 2013). The cytosol and stroma conductances are difficult to estimate but are expected to be large, so they likely play a small role in determining $g_m$ and can be neglected (Ellsworth et al. 2018). Under these assumptions, the mesophyll conductance can be expressed as

$$\frac{1}{g_m} = \frac{1}{g_{ias}} + \frac{1}{g_{cw} \cdot S_c} + \frac{1}{g_{mem} \cdot S_c}, \quad (1)$$

where $S_c$ is the chloroplast surface area appressing intercellular airspace per unit leaf area ($m^2$ chloroplast $m^{-2}$ leaf), $g_m$ and $g_{ias}$ are expressed on a leaf area basis (mol $m^{-2}$ leaf $s^{-1}$ $bar^{-1}$), $g_{cw}$ and $g_{mem}$ are expressed on a chloroplast area basis (mol $m^{-2}$ chloroplast $s^{-1}$ $bar^{-1}$), and $\frac{1}{g_{mem}} = \frac{1}{g_{pm}} + \frac{1}{g_{clm}}$. The factor $S_c$ converts conductances from a chloroplast area basis to a leaf area basis.

The cell wall conductance is determined from its anatomy according to

$$g_{cw} = \frac{p \cdot D_{liq} \cdot K_{CO_2}}{\tau \cdot T_{cw}} \cdot \frac{1}{R \cdot T_{leaf}}, \quad (2)$$

where $p$ is the cell wall porosity (dimensionless), $D_{liq}$ is the diffusivity of $CO_2$ in water ($m^2$ $s^{-1}$), $K_{CO2}$ is the partition coefficient for $CO_2$ in water (dimensionless), τ is the tortuosity of the $CO_2$ diffusion path through the cell wall (dimensionless), $T_{cw}$ is the cell wall thickness (m), $T_{leaf}$ is the leaf temperature (K), and $R$ is the ideal gas constant (J $k^{-1}$ $mol^{-1}$) (Ellsworth et al. 2018). Dividing by $R \cdot T_{leaf}$ converts the conductance from m $s^{-1}$ to mol $m^{-2}$ $s^{-1}$ bar as discussed below. Porosity and tortuosity are difficult to estimate, and can instead be combined into an effective porosity $p / \tau$ (Ellsworth et al. 2018). Combining Equations 1 and 2, the effective porosity can be calculated as

$$\frac{p}{\tau} = \frac{T_{cw} \cdot R \cdot T_{leaf} \cdot g_{cw}}{D_{liq} \cdot K_{CO_2}} = \frac{T_{cw} \cdot R \cdot T_{leaf}}{D_{liq} \cdot K_{CO_2}} \cdot \left[\frac{S_c}{g_m} - \frac{S_c}{g_{ias}} - \frac{1}{g_{mem}}\right]^{-1}. \quad (3)$$

This is a useful expression because all factors on the right-hand-side can either be measured or calculated, enabling an estimate of the effective porosity. The conductance in the intercellular airspace can be calculated as

$$g_{ias} = \frac{D_{air} \cdot f_{ias}}{0.5 \cdot T_{mes} \cdot \xi} \cdot \frac{1}{R \cdot T_{leaf}}, \quad (4)$$

where $D_{air}$ is the diffusivity of $CO_2$ in air, $f_{ias}$ is the fraction of intercellular airspace within the mesophyll, $T_{mes}$ is the mesophyll thickness, and $\xi$ is the tortuosity of the $CO_2$ diffusion path through the intercellular airspace (Xiong 2023). The membrane conductance can be calculated as

$$g_{mem} = (1 + \epsilon_{mem}) \cdot g_{mem}^{bilayer}, \quad (5)$$

where $g_{mem}^{bilayer}$ is the membrane conductance as estimated from the permeability of a lipid bilayer (given by Equation 5 in (von Caemmerer and Evans (2015)) and $\epsilon_{mem}$ is a dimensionless conductance enhancement factor accounting for facilitation processes (Xiong 2023). Here we use $\epsilon_{mem} = 2$, as estimated from the Solanaceae crops (potato and tomato) included in Figure 7d of Xiong (2023).

The diffusivity of $CO_2$ in water ($D_{liq}$) can be calculated using Equation 4 from von Caemmerer and Evans (2015). The cell wall thickness $T_{cw}$, mesophyll thickness $T_{mes}$, and fraction of intercellular airspace $f_{ias}$ can be determined from microscopy images. The mesophyll conductance $g_m$ can be estimated from combined gas exchange and isotope discrimination measurements. The leaf temperature (25 °C) is known from measurement conditions, and temperatures are converted from °C to K by adding 273 (von Caemmerer and Evans 2015). The values of the remaining parameters were:

- $K_{CO2}$ = 1 (Nobel 2009)
- $R$ = 8.314 J $K^{-1}$ $mol^{-1}$ (von Caemmerer and Evans 2015)
- $D_{air}$ = 1.515 x $10^{-5}$ $m^2$ $s^{-1}$ (Xiong 2023)
- $\xi$ = 1.57 m $m^{-1}$ (Xiong 2023)

Care must be taken when calculating $g_{cw}$ and $g_{ias}$ from anatomical parameters. The equations as written in the sources cited calculate these conductances in units of m $s^{-1}$. Thus, they are intended to be used along with gas concentrations expressed in vapor densities (kg $m^{-3}$) rather than partial pressures (bar). It can be shown using the ideal gas law that conductances in m $s^{-1}$ can be divided by $R \cdot T_{leaf}$ to convert to the correct units, hence the appearance of this factor in our Equations 2 and 4. (Note that 1 J = 1 Pa $m^3$ and 1 bar = $10^5$ Pa.)

Equation 3 for the effective porosity is similar to the approach taken in Ellsworth et al. (2018), but it also includes the conductance through the intercellular airspaces and a membrane conductance enhancement factor as in Xiong (2023).

**Supplementary Dataset 1. Codon optimized sequence of AtCGR3**

'AtCGR3' is original Arabidopsis CGR3 full length coding sequence (AT5G65810.1)

'AtCGR3_NtCO' is Arabidopsis CGR3 sequence codon optimized for *Nicotiana tabacum*

'.' indicates base remains unchanged

```
Identities:581/777(75%), Strand: Plus/Plus

AtCGR3       1    ATGTCAAGAAGGCAAGTAAGGCGTGTAGGGGATAGTGGAAGCTTCCCATTTGTAGGAGCT  60
AtCGR3_NtCO  1    ..............G..C...........A..CTCC..GTCA.....T..C..T......  60

AtCGR3       61   CTGCATTCAAAATCACGTTCGTCTCCTCTGTTATCAGTTTGCCTTGTTCTCGTGGGAGCA  120
AtCGR3_NtCO  61   T.A..CAGC.....T.....C........C.....C..C..TT.A..A.....T..T..C  120

AtCGR3       121  TGCCTTCTCATTGGTTATGCTTACAGTGGTCCAGGTATGTTCAAAAGTATCAGAGAAGTC  180
AtCGR3_NtCO  121  .....CT.G........C..C...TC...A..T...........G..C.....G.....T  180

AtCGR3       181  AGCAAGATTACAGGTGACTATTCTTGCACAGCAGAAGTTCAAAGAGCCATTCCTATTCTT  240
AtCGR3_NtCO  181  ...........T..C..T.....A.....C..T..G..A...C.T...........T.G  240

AtCGR3       241  AAGAGTGCGTATGGAGATAGCATGCGCAAAGTCCTGCACGTGGGTCCTGAAACATGCTCA  300
AtCGR3_NtCO  241  ..A..C..T.....T...TCA.....A..G..TT....T..T..G.....G..T......  300

AtCGR3       301  GTGGTCTCGAGTCTGTTGAATGAAGAAGAGACAGAAGCATGGGGTGTTGAACCATATGAT  360
AtCGR3_NtCO  301  ..A...AGCTCA...C....C..G...........G..T.....C..A.....T..C...  360

AtCGR3       361  GTGGAGGATGCAGACTCTAACTGCAAAAGTCTTTTGCACAAGGGCCTTGTACGTGTGGCT  420
AtCGR3_NtCO  361  ..C..A...........C..T.....GTCA..G.....T..A..T..G.....A..C..A  420

AtCGR3       421  GACATCAAATTCCCTCTTCCTTACCGGTCAAAGTCGTTTTCTCTTGTGATCGTCTCAGAC  480
AtCGR3_NtCO  421  ........G..T..CT.G..A..TA....T.....A........C...........C..T  480

AtCGR3       481  GCTTTGGATTACCTCTCACCCAGGTACCTGAACAAAACTGTGCCTGAACTTGCTCGCGTC  540
AtCGR3_NtCO  481  ..AC.C........G........A...T.A..T..G........C.....G.....T..T  540

AtCGR3       541  GCTTCAGATGGTGTCGTTCTTTTAGCAGGTAACCCTGGTCAACAAAAGGCTAAAGGTGGG  600
AtCGR3_NtCO  541  ..AAGT........A..C...C.T..T..A.....A..A........A.....G.....T  600

AtCGR3       601  GAATTGTCGAAATTTGGACGGCCTGCTAAAATGCGTAGCTCGTCG-TGGTGGATCCGTTT  659
AtCGR3_NtCO  601  ......AGT.........A....C.........A.G...-A..A.C........AA.A..  659

AtCGR3       660  CTTCTCACAGACGAACTTAGAGGAAAACGAAGCAGCAAGCAAGAAATTCGAACAAGCAGC  719
AtCGR3_NtCO  660  T..TAGC.....A...C.C..A..G..T.....C...TC.........T..G........  719

AtCGR3       720  TTCCAAGAGTTCATACAAACCAGCTTGTCAAGTTTTCCACCTCAAGCCATTACATTAG  777
AtCGR3_NtCO  720  CAGT...TCC..C.....G..T..A........G..T..TT.G.....T.....CGGT  777
```
